# Frontal noradrenergic and cholinergic transients exhibit distinct spatiotemporal dynamics during competitive decision-making

**DOI:** 10.1101/2024.01.23.576893

**Authors:** Hongli Wang, Heather K. Ortega, Emma B. Kelly, Jonathan Indajang, Jiesi Feng, Yulong Li, Alex C. Kwan

**Author notes:** These authors contributed equally to the work (Hongli Wang, Heather K. Ortega). **Author Contributions:** H.W., H.K.O., and A.C.K. designed research; H.W., H.K.O., E.B.K., and J.I. performed research. H.W., H.K.O., and A.C.K. analyzed data. J.F. and Y.L. contributed new reagents. H.W. and H.K.O. wrote the first draft of the paper; H.W., H.K.O., E.B.K., J.I., and A.C.K. edited the paper; H.W., H.K.O., and A.C.K. wrote the paper.

## Abstract

Norepinephrine (NE) and acetylcholine (ACh) are neuromodulators that are crucial for learning and decision-making. In the cortex, NE and ACh are released at specific sites along neuromodulatory axons, which would constrain their spatiotemporal dynamics at the subcellular scale. However, how the fluctuating patterns of NE and ACh signaling may be linked to behavioral events is unknown. Here, leveraging genetically encoded NE and ACh indicators, we use two-photon microscopy to visualize neuromodulatory signals in the superficial layer of the mouse medial frontal cortex during decision-making. Head-fixed mice engage in a competitive game called matching pennies against a computer opponent. We show that both NE and ACh transients carry information about decision-related variables including choice, outcome, and reinforcer. However, the two neuromodulators differ in their spatiotemporal pattern of task-related activation. Spatially, NE signals are more segregated with choice and outcome encoded at distinct locations, whereas ACh signals can multiplex and reflect different behavioral correlates at the same site. Temporally, task-driven NE transients were more synchronized and peaked earlier than ACh transients. To test functional relevance, using optogenetics we found that evoked elevation of NE, but not ACh, in the medial frontal cortex increases the propensity of the animals to switch and explore alternate options. Taken together, the results reveal distinct spatiotemporal patterns of rapid ACh and NE transients at the subcellular scale during decision-making in mice, which may endow these neuromodulators with different ways to impact neural plasticity to mediate learning and adaptive behavior.

## Introduction

Neuromodulators including acetylcholine (ACh) and norepinephrine (NE) play pivotal roles in various behavioral functions (Everitt and Robbins, 1997; Aston-Jones and Cohen, 2005; Picciotto et al., 2012). One function associated with central cholinergic tone is arousal and vigilance (Buzsaki et al., 1988), which relate to sensory sensitivity and selective attention (Dalley et al., 2001; Yu and Dayan, 2002). These functions are supported by many experiments that manipulated cholinergic signaling using pharmacology, lesions, and optogenetics (McGaughy et al., 2002; Chudasama et al., 2004; Parikh et al., 2007; Herrero et al., 2008; Pinto et al., 2013; Gritton et al., 2016). Classically, perceptual effects are associated with slow fluctuation of ACh levels, although recent evidence indicates control can also occur at more rapid timing (Parikh et al., 2007; Goard and Dan, 2009). Relatedly, NE has also been implicated in arousal and vigilance (Jouvet, 1969; McCormick et al., 1991), and improved sensitivity to sensory cues (Berridge and Waterhouse, 2003). This is possibly achieved by NE elevating the signal-to-noise ratio and/or gain in neural networks (Servan-Schreiber et al., 1990; Eldar et al., 2013). The neuromodulatory effects on behavior, such as those exerted by NE, exhibit an inverted U-shaped curve (Aston-Jones et al., 1999). Indeed, activity of cortical cholinergic and noradrenergic axons correlate well with pupil diameter, which is an indicator of the arousal level of an animal (Reimer et al., 2016).

In addition to arousal and vigilance, it is established that ACh and NE may be important for learning and decision-making (Hasselmo and Bower, 1993; Doya, 2002; Bouret and Sara, 2005; Dayan and Yu, 2006; Sara, 2009). For instance, cholinergic neurons in the basal forebrain exhibit fast and transient increase in spiking activity after primary reinforcements including rewards and punishments (Lin and Nicolelis, 2008; Hangya et al., 2015). In support, optogenetic activation of cortical cholinergic axons could substitute for actual rewards in associative learning (Liu et al., 2015). Memory deficits have been observed when cholinergic transmission was abolished in animals (Chudasama et al., 2004; Croxson et al., 2011). NE may be similarly crucial for learning and decision-making because a loss of adrenergic receptors in the prefrontal cortex contributes to the memory loss in aged animals (Arnsten and Goldman-Rakic, 1985; Arnsten et al., 2012). More specifically, locus coeruleus neurons fire at specific epochs during decision tasks (Clayton et al., 2004) and may be sensitive to reward values (Bouret and Richmond, 2015). Presumably, neuronal firing changes in cholinergic or noradrenergic nuclei reflect the altered phasic release of these neuromodulators that have been observed in the cortex (Teles-Grilo Ruivo et al., 2017), which may drive long-term synaptic plasticity in cortical circuits (Kilgard, 1998; Froemke et al., 2007; Martins and Froemke, 2015). Therefore, growing evidence indicate functions of ACh and NE signaling in higher cognitive functions.

Despite the large body of literature showing that ACh and NE can have multiple behavioral functions and act at multiple timescales, less is known about the spatial pattern of the neuromodulatory signal. At the level of brain regions, the neuromodulatory signals come from different sources and have distinct projection patterns. The major source of ACh in the neocortex comes from the basal forebrain. The axonal projections are organized topographically (Saper, 1984; Zaborszky et al., 2015; Kim et al., 2016), and exhibit rich spatiotemporal dynamics across regions (Lohani et al., 2022). By contrast, a major source of NE is locus coeruleus, which sends axons to innervate much of the forebrain – though with some exceptions, such as the basal ganglia (Amaral and Sinnamon, 1977; Moore and Bloom, 1979). Unlike the cholinergic system, each locus coeruleus neuron projects broadly to many brain regions (Loughlin et al., 1982; Schwarz et al., 2015) with a high divergence of >20,000 terminals (Descarries and Lapierre, 1973). At a finer spatial scale, within a single brain region, ACh and NE signaling must be heterogeneous in space because neuromodulator levels elevate at locations of release sites of the respective neuromodulatory axonal fiber terminals (Zhu et al., 2020). However, the extent to which the spatial patterns of ACh and NE signals at the subcellular resolution may reflect behavioral events is unknown.

In this study, we address this knowledge gap by leveraging the latest generation of genetically encoded fluorescent indicators of ACh (Jing et al., 2020) and NE (Feng et al., 2019; Feng et al., 2023), which permit sensitive and spatially resolved imaging of neuromodulatory signals. We trained head-fixed mice to play a competitive game called matching pennies against a computer opponent (Wang et al., 2022). In the medial frontal cortex, we found that although both NE and ACh transients encoded the same set of task-related variables on a trial-by-trial basis, their spatiotemporal dynamics are different. NE at a location would encode often only one decision variable, whereas ACh at one site tends to multiplex and be driven by different behavioral events. To determine behavioral relevance, we activate cholinergic or noradrenergic fibers in the medial frontal cortex using optogenetics to show that increased NE availability selectively promotes exploration during decision-making.

## Results

### Head-fixed mice play matching pennies against a computer opponent

Matching pennies is a competitive game that involves social and strategic decision-making (Lee, 2008; Wang and Kwan, 2023). We previously developed a behavioral paradigm for head-fixed, fluid-restricted mice to play matching pennies against a computer opponent and characterized the behavioral performance in detail (Wang et al., 2022). Briefly, in this iterated version of matching pennies, for each trial, the animal and the computer chose simultaneously the left or right option (**Fig. 1A**). Outcome was determined by a payoff matrix: if the mouse chose the same option as the computer, the mouse received a water reward; otherwise, there was no reward (**Fig. 1B**). The computer opponent was programmed to predict the animal’s upcoming choice using the choice and outcome history over the session (see **Methods**). Based on the prediction, the computer aimed to provide competitive pressure by selecting the option that the mouse is less likely to pick. At the beginning of each trial, a 0.2-s, 5-kHz sound cue was played to initiate a 2-s response window, during which the mouse could indicate its choice by licking either the left or right spout with its tongue (**Fig. 1C**). Based on the choices of the animal and the computer, a water reward might be delivered at the chosen spout according to the payoff matrix. A random intertrial interval was presented to suppress pre-cue licks, which would be prolonged if the animal emitted one or more licks during the interval (see **Methods**).

**Figure 1.**
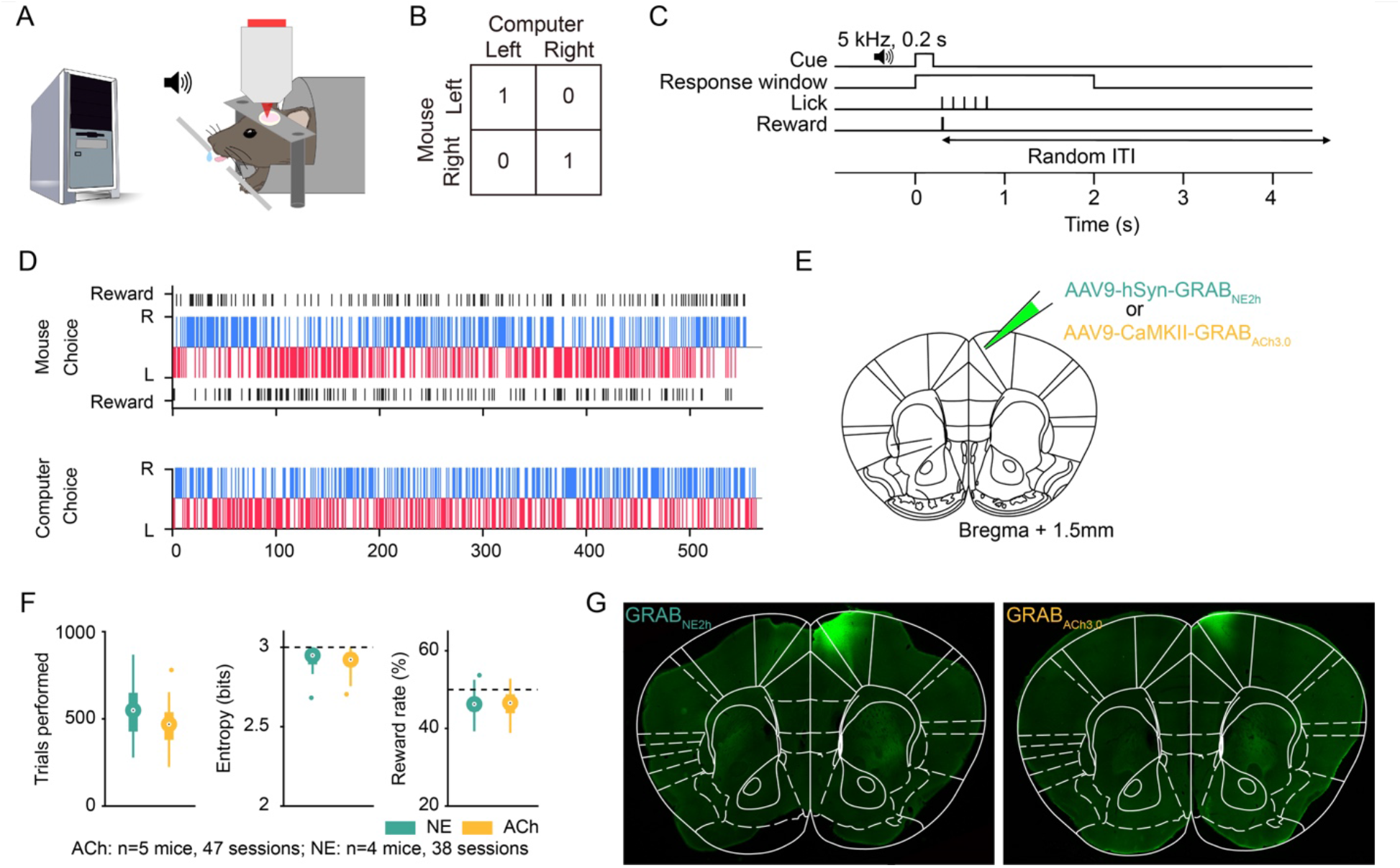
Head-fixed mice play matching pennies against a computer opponent. **(A)** Schematic of the competitive game. Head-fixed mouse licks left or right spout to indicate left or right choices. A computer tracks the animal’s previous choices and outcomes, and chooses the side that the mouse is less likely to pick. **(B)** The payoff matrix of the game. The mouse receives a water reward only if it and the computer choose the same action in a trial. **(C)** Each trial, a sound cue signals the start of a response window. The first lick emitted by the animal within the window is logged as the response for that trial, and the outcome is delivered immediately based on the payoff matrix. A random intertrial interval follows the outcome. **(D)** An example session in which the mouse performed at a 50.3% reward rate. Top: the mouse’s choices and outcomes. Bottom: the computer’s choices. Blue and red bars indicate right and left choices, respectively. Black bars indicate rewards. **(E)** Schematic of the injection site. **(F)** Summary from 38 GRAB_NE2h_ (green) and 47 GRAB_ACh3.0_ (yellow) sessions. Left: the average number of trials performed each session. Middle: the average entropy of 3-choice sequences. Right: the average reward rate. **(G)** Post hoc widefield fluorescence image of GRAB_NE2h_- and GRAB_ACh3.0_-expressing neurons, immunostained with an anti-GFP antibody, in the medial M2 region of the frontal cortex.

In preparation for characterizing noradrenergic and cholinergic transients, adult C57BL/6J mice were injected with AAV9-hSyn-GRAB_NE2h_ or AAV9-CaMKII-GRAB_ACh3.0_ to express genetically encoded fluorescent indicators of NE (Feng et al., 2019; Feng et al., 2023) and ACh (Jing et al., 2020) in the medial secondary motor cortex (M2) region of the medial frontal cortex (**Fig. 1E**). We focused on the medial M2 region because of its role in flexible decision-making (Siniscalchi et al., 2016; Barthas and Kwan, 2017; Yang and Kwan, 2021). Headplate and cranial glass window were implanted to enable head fixation and cellular-resolution optical imaging. Animals were trained to reach a stable performance of >40% reward rate for 3 consecutive sessions. The Nash equilibrium of matching pennies suggests that the optimal play is a mixed strategy: players should choose left and right with equal probabilities, which would yield a 50% reward rate in the long run. Indeed, the animals made choices with a high degree of stochasticity in a session (**Fig. 1D**). In total, the data set involving two-photon imaging during matching pennies included 47 sessions from 5 animals expressing GRAB_ACh3.0_ and 38 sessions from 4 animals expressing GRAB_NE2h_. On average, animals expressing GRAB_NE2h_ and GRAB_ACh3.0_ sensors performed 550±25 and 459±17 trials per session respectively (mean±s.e.m.; **Fig. 1F**). Both groups exhibited a high level of stochasticity in choice behavior, exemplified by the mean entropy values of 2.93±0.01 and 2.91± 0.01 for the NE and ACh groups. Accordingly, the animals received reward rates of 46.2±0.5% and 46.2±0.5%, which were near but lower than the optimal reward rate of 50% (*p*=4.5×10^-10^ for NE; *p*=4.5×10^-10^ for ACh; Wilcoxon rank sum test). Post hoc histology showed the spatial extent of the GRAB_NE2h_ and GRAB_ACh3.0_ expression in the medial frontal cortex (**Fig. 1G**). Together, these results showed that animals undergoing two-photon imaging can play matching pennies at an expert level.

### Visualizing frontal cortical NE and ACh transients using genetically encoded fluorescent indicators

We used a two-photon microscope to record fluorescence signals from GRAB_NE2h_ and GRAB_ACh3.0_ sensors at a depth of 100-150 µm below the dura (**Fig. 2A**). The fluorescence signals were diffuse across the field of view (**Fig. 2B, C**), presumably because the sensors express densely in cell bodies as well as neuropil. There were typically several dark areas in a field of view, which were likely blood vessels and capillaries. Because of the diffuse signal, we decided to divide the field of view in an unbiased manner by using an evenly spaced grid of 28×28 regions of interest (ROI). Each ROI had an area of 4.46×4.46 μm. We tested several coarser and finer grid spacings, and found that it did not affect the conclusions of the subsequent analyses.

**Figure 2.**
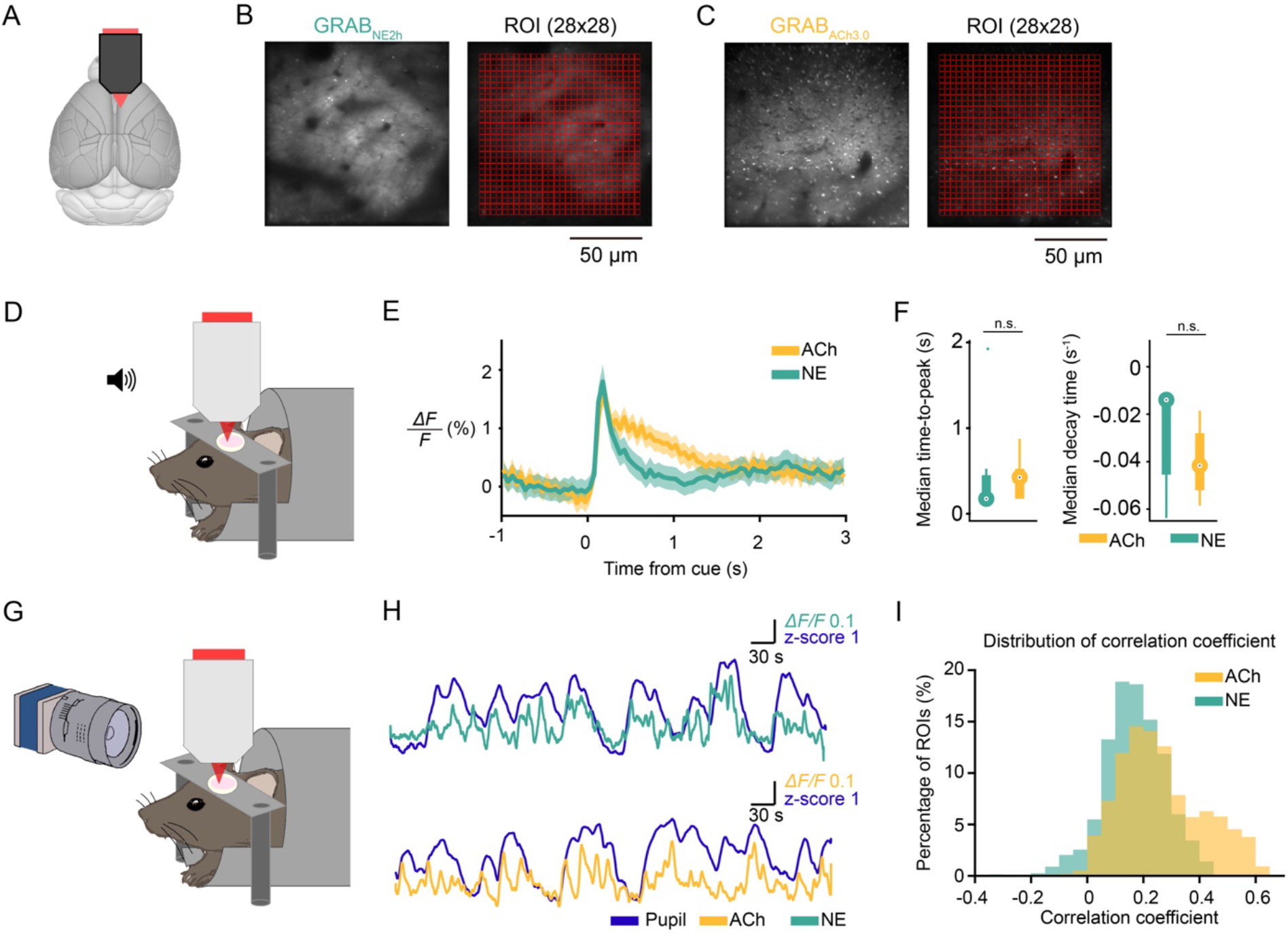
Visualizing frontal cortical NE and ACh transients using genetically encoded fluorescent indicators. **(A)** Schematic of the imaging site. **(B)** Example field of view of GRAB_NE2h_-expression in layer 2/3 of M2 imaged *in vivo* with two-photon microscopy. A 28x28 grid was used to divide the field of view into regions of interest (ROIs). **(C)** Similar to (B) for GRAB_ACh3.0_. **(D)** Schematic of setup to record auditory evoked response. A 4-kHz, 50-ms auditory stimulus was presented 200 times per session, with simultaneous two-photon imaging. **(E)** Auditory evoked responses averaged across all ROIs and sessions for GRAB_NE2h_ (green) and GRAB_ACh3.0_ (yellow). Shading represents the 95% confidence interval. **(F)** Temporal parameters of the auditory evoked responses. Left: median time-to-peak from cue time. GRAB_ACh_: 0.43 (0.18 – 0.53) s; GRAB_NE_: 0.18 (0.18 – 0.45) s. Data are reported as median (25th – 75th percentile). *p* = 0.7, Wilcoxon rank sum test. Right: median decay time. GRAB_NE:_ -0.01 (-0.04 – -0.01) s_-1_; GRAB_ACh_: -0.04 (-0.05 – -0.03) s_-1_. *p* = 0.16, Wilcoxon rank sum test. n.s., not significant. Circle, median. Thick bar, 25th and 75th percentiles. Thin bar, maximum and minimum values. **(G)** Schematic of setup to record pupil diameter. Pupil and two-photon images were recorded simultaneously. **(H)** Spontaneous pupil diameter in z-score (purple) and GRAB_NE2h_ (green) or GRAB_ACh3.0_ (yellow) signal from an example ROI. **(I)** Distributions of maximal coefficients in cross-correlation between pupil dilations and GRAB_NE2h_ (green) or GRAB_ACh3.0_ (yellow) signal.

We performed two experiments to confirmed that the fluorescence signals arising from GRAB_NE2h_ and GRAB_ACh3.0_ sensors reflect NE and ACh transients in the medial frontal cortex. One, we recorded auditory evoked response, because previous studies showed that both noradrenergic neurons in LC and cholinergic neurons in basal forebrain respond to auditory stimuli, particularly novel and unexpected cues (Moore and Bloom, 1979; Maho et al., 1995). Indeed, presentation of 4-kHz, 50-ms auditory cues led to a sharp-rising fluorescent transient from animals that expressed GRAB_NE2h_ or GRAB_ACh3.0_ sensor (**Fig. 2D-F**). Two, cortical cholinergic and noradrenergic axonal activities are correlated with pupillary fluctuations (Reimer et al., 2016). We measured spontaneous fluctuations in pupil diameter while imaging GRAB_NE2h_ and GRAB_ACh3.0_ signals in the medial frontal cortex (**Fig. 2G**). As expected, periods of pupil dilation corresponded roughly to periods of elevated fluorescence signals (**Fig. 2H**). Collating data from all ROIs across all fields of view, pupil size and fluorescence signals were positively correlated in most ROIs for both GRAB_NE2h_ and GRAB_ACh3.0_ (**Fig. 2I**). These results provided evidence that the fluorescence signals from GRAB_NE2h_ and GRAB_ACh3.0_ sensors acquired with two-photon microscopy reported fluctuations of NE and ACh levels in the medial frontal cortex.

### Frontal cortical NE and ACh transients contain choice- and outcome-related signals

We imaged fluorescence signals to determine NE and ACh transients in the medial frontal cortex as mice played matching pennies against a computer opponent (**Fig. 3A**). We observed NE and ACh transients that differed for rewarded versus unrewarded trials and contralateral versus ipsilateral choices (**Fig. 3B**). To determine more quantitatively how NE and ACh transients in all ROIs relate to behavioral events, we fitted a multiple linear regression model (see **Methods**) for each ROI to determine how its fluorescence signal may be explained by choices, outcomes, and reinforcers (choice-outcome interactions) of past two, current, and next trials as well as recent reward rate and cumulative reward sum in a session. The results revealed that NE and ACh transients in a sizable fraction of ROIs were modulated by choice, outcome, and reinforcer of the current trial (**Fig. 3C**). We noted several differences between NE and ACh. For choice, the ACh signal rose before the cue, while it was detected in NE only after the cue onset. This is consistent with our earlier finding that pupil-related arousal contained choice information prior to cue (Wang et al., 2022) and indicates that cortical ACh, but not NE, may be involved in the preparation of the upcoming action. There were also differences in the temporal profiles of the outcome-related ACh and NE signals, which will be examined quantitatively in the next sections. To a lesser degree, frontal cortical ACh and NE levels were modulated by other behavioral predictors including previous choice, previous outcome, recent reward rate, and cumulative reward sum (**Fig. 3D**). This analysis showed that NE and ACh transients in the medial frontal cortex vary with decisions during the competitive game.

**Figure 3.**
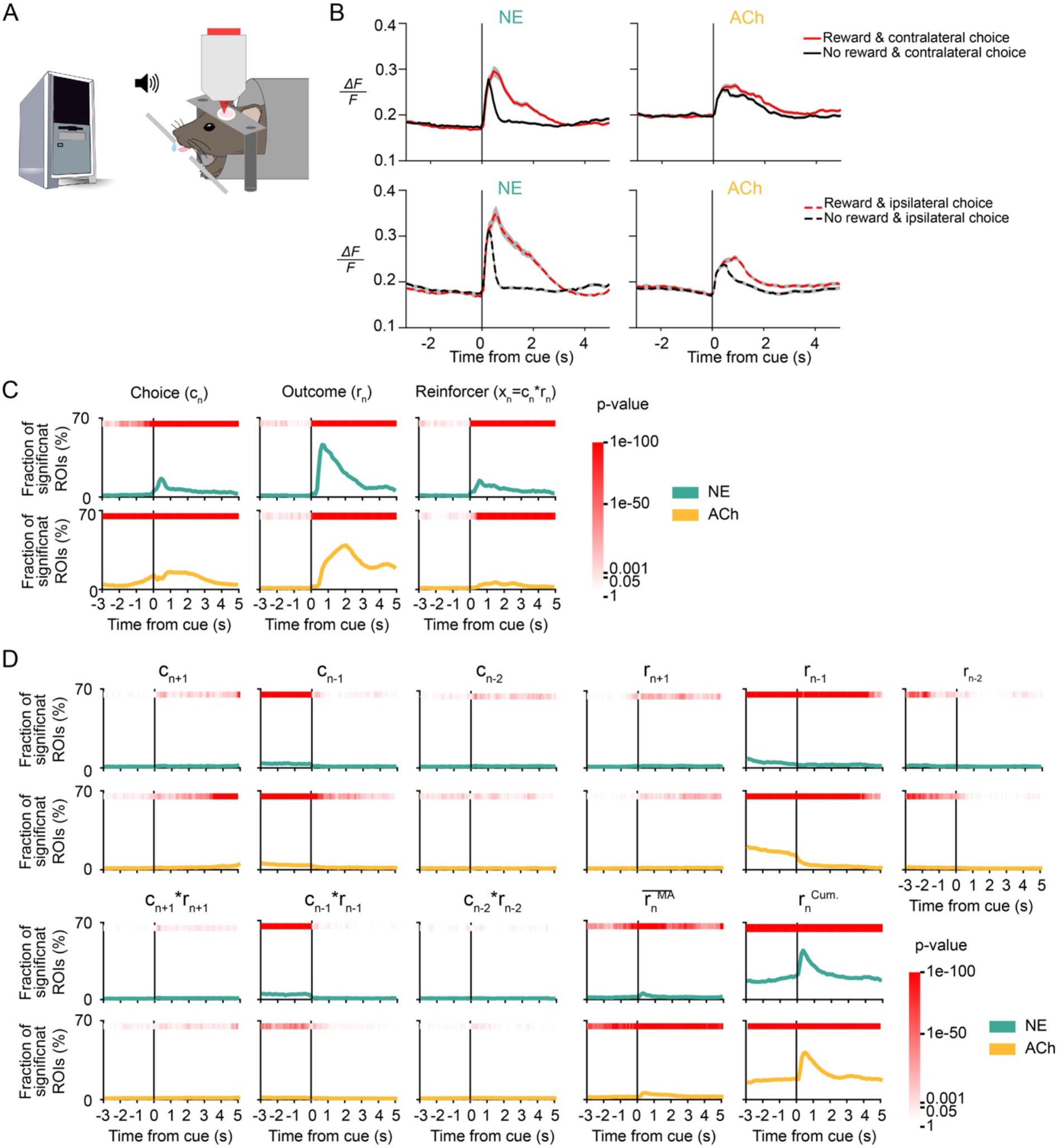
Frontal cortical NE and ACh transients contain choice- and outcome-related signals. **(A)** Schematic of the mouse playing matching pennies while NE or ACh transients were imaged using two-photon microscopy. **(B)** Trial-averaged fluorescence traces aligned to the cue onset for different subsets of trials. One example ROI was shown each for GRAB_NE2h_ (left) and GRAB_ACh3.0_ (right). Gray shading: 95% confidence interval. **(C)** The proportion of ROIs with significant regression coefficient for choice in the current trial *c_n_*, outcome in the current trial *r_n_*, and reinforcer (choice-outcome interaction) in the current trial *x_n_* in GRAB_NE2h_ (green) and GRAB_ACh3.0_ (yellow) data, determined by fitting a multiple linear regression model. Red shading indicates the p-value from the chi-square test. **(D)** The fraction of ROIs with significant regression coefficient for choice in the next trial *c_n+1_*, choice in the previous trial *c_n-1_*, choice in the trial before the previous trial *c_n-2,_* outcome in the next trial *r_n+1_*, outcome in the previous trial *r_n-1_*, outcome in the trial before the previous trial *r_n-2,_* reinforcer in the next trial *x_n+1_*, reinforcer in the previous trial *x_n-1_*, reinforcer in the trial before the previous trial *x_n-2,_* recent reward rate *r*_n_ ^MA^, calculated as a moving average over last 20 trials, and the cumulative reward *r*_n_^cum^ from start of session to current trial for GRAB_NE2h_ (green) and GRAB_ACh3.0_ (yellow) data determined from the same fit as (B). Red shading indicates the *p*-value from the chi-square test.

### Spatial organization of the decision-related NE and ACh transients

A key advantage of two-photon imaging is to obtain micron-scale maps of the ACh and NE signals (**Fig. 4A**). We can average coefficients obtained from multiple linear regression across time and ask how fluorescent transients at different subcellular locations in the medial frontal cortex were linked to behavioral variables (**Fig. 4B**). **Fig. 4C** shows such analysis applied to one field of view, revealing heterogeneity in the spatial distribution of the task representations in the ACh and NE fluctuations. Across all sessions, we found that 81.9% and 83.3% of the ROIs had GRAB_NE2h_ and GRAB_ACh3.0_ transients that were modulated by at least one of the behavioral variables in the multiple linear regression model (**Fig. 4D**). Focusing on the choice, outcome, and reinforcer in the current trial that constitute the most predictive behavioral variables, 19.2% and 26.9% of the NE and ACh ROIs were significantly modulated by choice (**Fig. 4E**). Meanwhile, more locations encoded outcomes, encompassing 46.6% and 60.6% of the ACh and NE ROIs. Finally, a minority of 18.0% and 9.9% of the NE and ACh ROIs were associated with reinforcer. A single ROI may be significantly modulated by more than one behavioral variable. This might happen because (1) locations represent behavioral variables with independent probabilities and overlap by chance or (2) locations may preferentially have correlated representation of multiple behavioral variables. The second explanation was supported by statistical tests, because the overlap of ROIs modulated by different behavioral variables occurred at a rate higher than chance for both ACh and NE (*p*=0, *p*=0, *p*=0, for overlap between choice- and outcome-, choice- and reinforcer-, and outcome-and reinforcer-modulated ROIs respectively, Pearson independent test; **Supplementary Table 4-1**).

**Figure 4.**
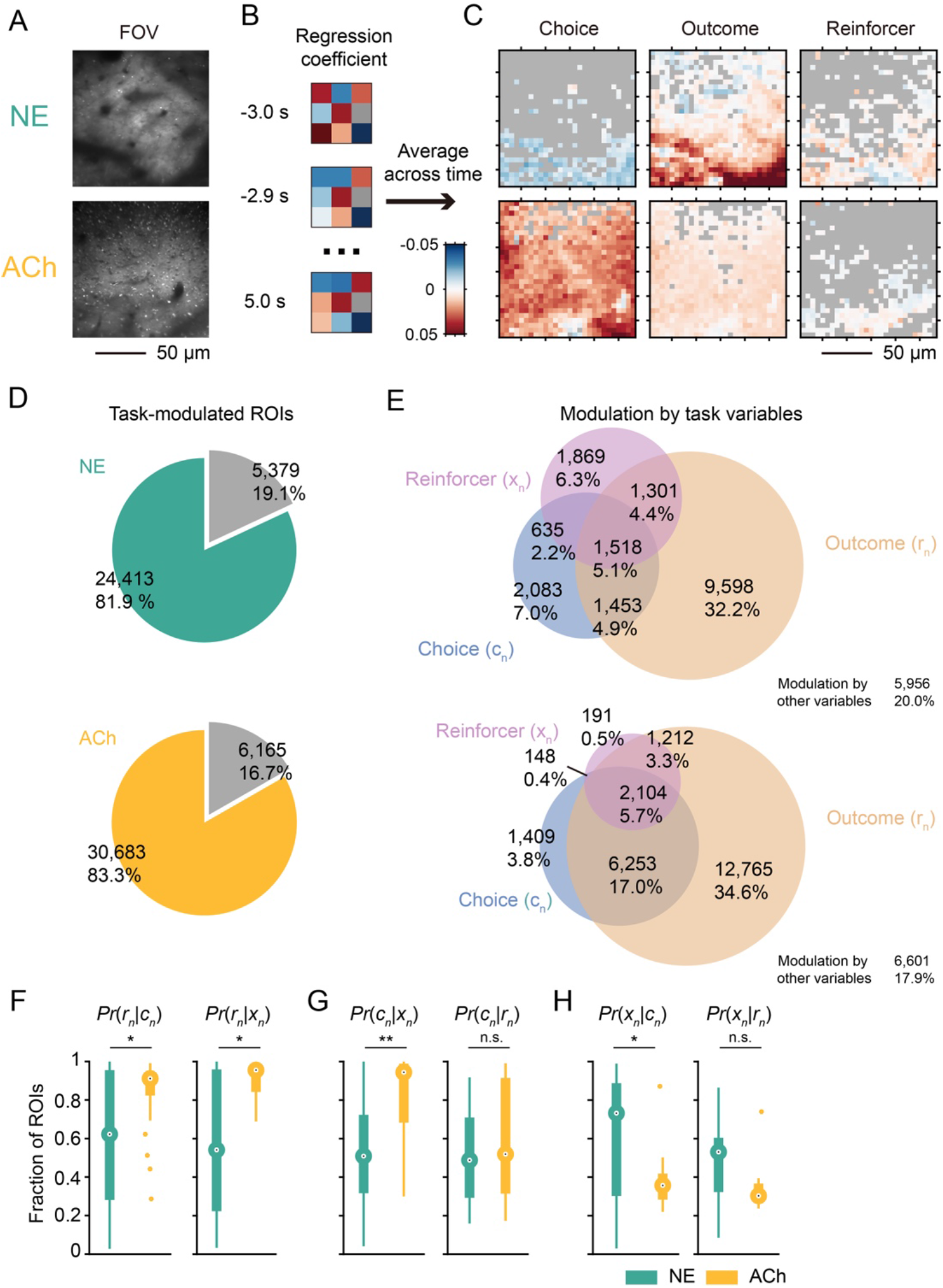
Spatial organization of the decision-related NE and ACh transients. **(A)** Example field of views of GRAB_NE2h_ (top) and GRAB_ACh3.0_ (bottom) expression in layer 2/3 of M2. **(B)** Schematic illustrating the analysis: for each ROI in a field of view, the regression coefficients over t = -3 - 5 s was averaged and plotted in pseudo-color. **(C)** Spatial maps of mean regression coefficients for choice (left), outcome (middle), and reinforcer (right) for the GRAB_NE2h_ (top row) and GRAB_ACh3.0_ (bottom row) fields of view shown in (A). **(D)** Venn diagrams showing the number and percentage of ROIs that were significantly modulated by at least one task-related variable. **(E)** Venn diagrams showing the number and percentage of ROIs that were significantly modulated by the current choice, outcome, and reinforcer. **(F)** Boxplot of the conditional probabilities in GRAB_NE2h_ (green) and GRAB_ACh3.0_ (yellow) data. Median test, *Pr*(*r_n_*|*c_n_*): *p*=0.044; *Pr*(*r_n_*|*x_n_*): *p*=0.034. \**p*<0.05; \*\**p*<0.01; n.s., not significant. Large circles, medians. Thick bars denote 25th and 75th percentiles. Lines end at maximum and minimum value. Small circles: outliers. **(G)** Same as (F) for *P_r_(c_n_|x_n_*): *p*=0.002; and *P_r_(c_n_|r_n_*): p = 0.864. **(H)** Same as (F) for *P_r_(x_n_|c_n_*): *p*=0.002; and *P_r_(x_n_|r_n_)*: *p* = 0.072.

To gain insight into the spatial integration of behaviorally relevant ACh and NE signals, we calculated the conditional probabilities that an ROI encoded one variable *v_1_* given that it also encoded another variable *v_2_* (*Pr*(*v_1_*|*v_2_*)). A head-to-head comparison of these conditional probabilities between NE and ACh highlighted a significantly higher degree of multiplexed coding of task information by ACh transients, as evident from the higher *Pr*(*r_n_*|*c_n_*), *Pr*(*r_n_*|*x_n_*), and *Pr*(*c_n_*|*x_n_*) values (*p*=0.044, *p*=0.034, and *p*=0.030; median test; **Fig. 4F, G; Supplementary Table 4-2)**. We did not detect a difference in *Pr*(*c_n_*|*r_n_*) (*p*=0.864; median test. **Fig. 4G)**. The *Pr*(*x_n_*|*c_n_*) and *Pr*(*x_n_*|*r_n_*) values were shown for completeness (*p*=0.002 and *p*=0.072; median test. **Fig. 4H**), but they represented a small number of ROIs due to fewer locations encoding reinforcer. Taken together, frontal cortical ACh transients are more likely to multiplex task-related information, where NE transients encode behavioral events in a more spatially segregated manner.

### Distinct temporal dynamics of the task-related NE and ACh signals

The most prominent behavioral readout linked to frontal cortical NE and ACh transients was outcome, therefore we asked how the reward-related signals evolve over time at different locations. To understand the spatiotemporal dynamics, we used the regression coefficient for outcome extracted for each ROI that was significantly modulated by outcome. These traces were sorted using hierarchical clustering based on Pearson correlation (**Fig. 5A**). The correlation matrices of the sorted regression coefficients revealed two clusters of ROIs for NE and ACh (**Fig. 5B**). For group 1, which captured 92.4% and 93.4% of the ROIs for NE and ACh respectively, the occurrence of a reward increased fluorescence signal (**Fig. 5C**). The temporal dynamics of NE and ACh signals differ, because NE signals rose faster and were more temporally aligned than ACh as reflected by the shorter median time-to-peak and smaller variance of time-to-peak (*p* = 1.54 × 10^-13^, and *p* = 0.002, respectively; Wilcoxon rank sum test; **Fig. 5D, left and middle**; **Supplementary Table 5-1**). The peak value of the regression coefficient was larger in NE than ACh (*p* = 0.003; Wilcoxon rank sum test; **Fig. 5F, right**; **Supplementary Table 5-1**), although this magnitude depended on experimental factors such as fluorophore expression level (Ali and Kwan, 2020) and therefore should be interpreted with caution. For group 2, which only captured 7.6% and 6.6% of the ROIs for NE and ACh respectively, the occurrence of a reward reduced fluorescence signal (**Fig. 3E**). There were similar differences in temporal dynamics for ROIs in group 2 as for those in group 1, except the difference in variance of time-to-peak was not statistically significant (median time-to-peak, *p* = 1.43 × 10^-4^; variance time-to-peak, *p* = 0.108; median peak-value, *p* = 0.002; Wilcoxon rank sum test; **Fig. 5F**; **Supplementary Table 5-1**).

**Figure 5.**
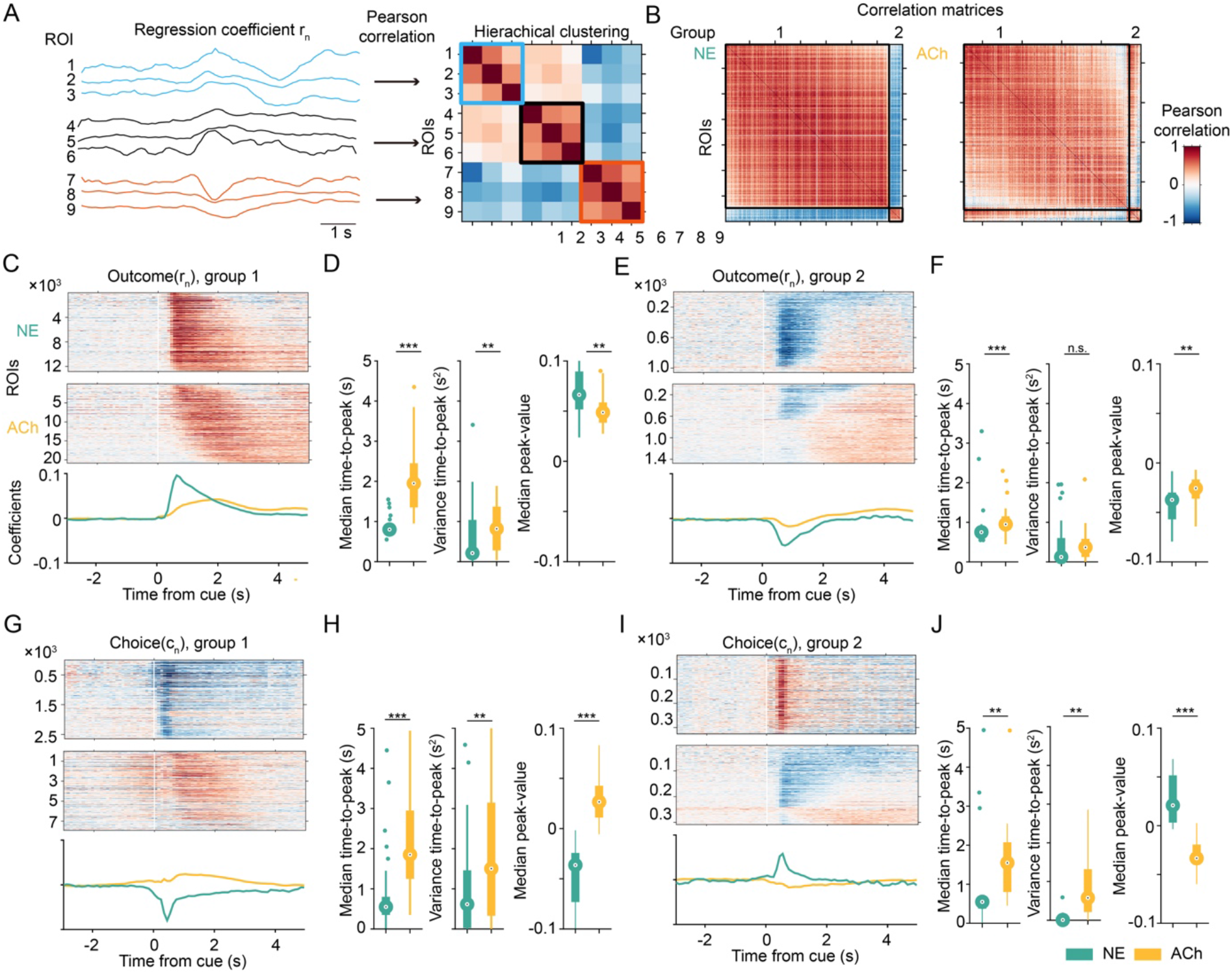
Distinct temporal dynamics of the task-related NE and ACh signals. **(A)** Schematic illustrating the analysis: regression coefficients for current outcome were clustered into different groups using hierarchical clustering based on Pearson correlation. **(B)** Correlation matrices showing the clustering results for GRAB_NE2h_ (left) and GRAB_ACh3.0_ (right). **(C)** Top: heat map of regression coefficients for current outcome for ROIs in group 1. Bottom: average regression coefficient for current outcome for ROIs in group 1 for GRAB_NE2h_ (green) and GRAB_ACh3.0_ (yellow). **(D)** Temporal parameters of outcome-related activity in group 1 for GRAB_NE2h_ (green) and GRAB_ACh3.0_ (yellow) data. Wilcoxon rank sum test. Left: median time-to-peak, *p*=1.54×10^-13^. Middle: variance time-to-peak, *p*=0.002. Right: median peak-value, *p*=0.003. \**p*<0.05; \*\**p*<0.01; \*\*\**p*<0.001; n.s., not significant. Large circle, median. Thick bar, 25th and 75th percentiles. Thin line, maximum and minimum values. Small circle, outlier. **(E, F)** Similar to (E) for ROIs in group 2. Median time-to-peak, *p*=1.43×10^-4^. Variance time-to-peak, *p*=0.108. Median peak-value, *p*=0.002. **(G, H)** Similar to (E) for regression coefficient for current choice for ROIs in group 1. Median time-to-peak, *p*=1.06×10^-7^. Variance time-to-peak, *p*=0.003. Median peak-value, *p*=3.53×10^-14^. **(I, J**) Similar to (E) for regression coefficient for current choice for ROIs in group 2. Median time-topeak, *p*=0.022. Variance time-to-peak, *p*=0.001. Median peak-value, *p*=1.88×10^-7^.

We clustered the ROIs based on their regression coefficient for outcome, but what about the choice-related signals, given that there is spatially correlated encoding of behavioral variables? We plotted the subset of ROIs with significant choice encoding using the same grouping and sorted ranking (**Fig. 5G, I**). The timing of the choice-related ACh and NE signals was also different: the choice-related NE signals emerged earlier and were more synchronized than ACh (group 1: median time-to-peak, *p*=1.06×10^-7^; variance time-to-peak, *p*=0.003; median peak-value, *p*=3.53×10^-14^; group 2: median time-to-peak, *p*=0.022; variance time-to-peak, *p*=0.001; median peak-value, *p*=1.88×10^-7^; Wilcoxon rank sum test; **Fig. 5H, J**; **Supplementary Table 5-1**). Strikingly, the sign of the regression coefficients for choice was opposite for NE and ACh. Although choice led to elevations of both NE and ACh in the medial frontal cortex, NE was preferentially driven by ipsilateral choice whereas ACh was more responsive to contralateral choice (**Fig. 5H**). These results reveal that decision-related NE signals were more synchronized and peaked earlier than ACh transients in the medial frontal cortex.

### Optogenetic elevation of frontal cortical NE increases switch probability

Given that NE and ACh transients exhibit task-related signals with distinct spatial and temporal dynamics, we wanted to know if their levels in the medial frontal cortex may differentially contribute to behavioral performance. To causally test the roles of the neuromodulators, we used optogenetics to stimulate noradrenergic and cholinergic axons in the medial frontal cortex as mice engaged in the matching pennies game (**Fig. 6A, B)**. The payoff matrix and trial structure were the same as the one used in imaging experiments except for the additional laser photostimulation on select trials **(Fig. 6C)**. Photostimulation (473 nm, 40 Hz) would start at cue onset and sustain until 1 s after the mouse makes a choice (i.e., the first lick within the response window). This was designed to roughly mimic the time course of the observed NE and ACh transients in the medial frontal cortex. We used a laser steering system that can rapidly re-position the laser beam and mice were implanted with a clear-skull cap (see **Methods**), which allowed us to photostimulate a different region in the dorsal cortex on each trial. To target noradrenergic neurons, we crossed a knock-in Dbh-Cre mouse (Tillage et al., 2020) with the Ai32 strain (Madisen et al., 2012) for Cre-dependent expression of ChR2. To target noradrenergic neurons, we crossed a ChAT-Cre mouse (Rossi et al., 2011) with the Ai32 strain for Cre-dependent expression of ChR2. Post hoc immunostaining and confocal microscopy of fixed coronal sections confirmed ChR2 expression in axons in the mouse medial frontal cortex (**Fig. 6D**).

**Figure 6.**
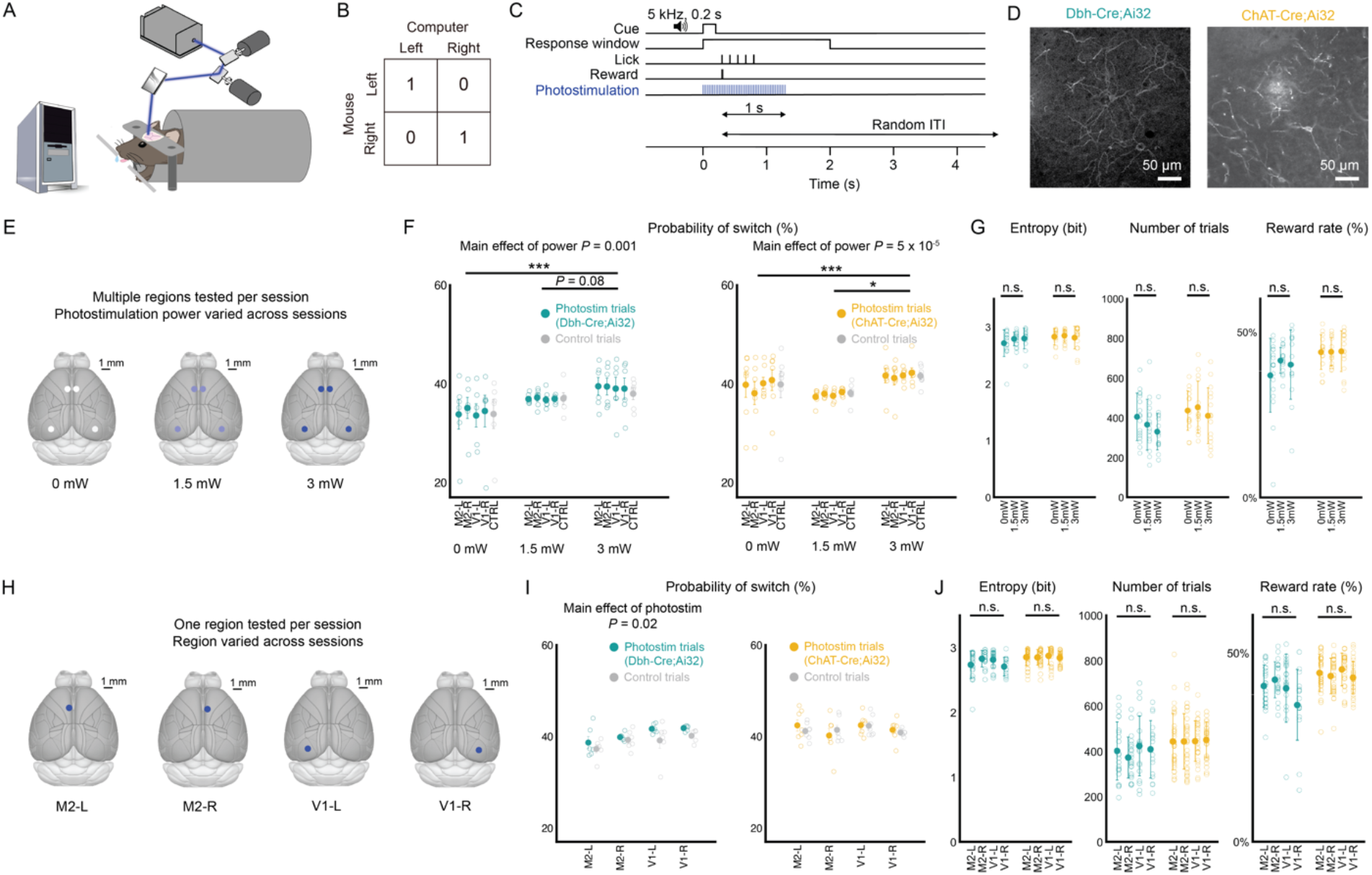
Optogenetic elevation of frontal cortical NE increases switching in matching pennies. **(A)** Schematic of the mouse playing matching pennies while noradrenergic and cholinergic axons were stimulated via optogenetics using a laser-steering system. **(B)** The payoff matrix of the game. **(C)** The timing of each trial. On trials with photostimulation, laser turns on at cue onset and sustains until 1 s after choice. **(D)** Confocal micrographs of immunostained axons in coronal section of medial frontal cortex of Dbh-Cre;Ai32 (left) and ChAt- Cre;Ai32 (right) mice. **(E)** Schematic of protocol to test effects of region within session and power across sessions. Multiple regions were tested per session, with maximum of one region tested per trial. 40% of trials are stimulated with 10% allocated to each region (M2-L, M2-R, V1-L, V1-R). One power level is tested per session. **(F)** Probability of switch on trial after photostimulation by region and power. **(G)** Session-based performance metrics, including entropy (left), number of trials (middle) and reward rate (right). **(H)** Schematic of protocol to test effects of regionspecific photostimulation across sessions. One region was tested per session. 10% of trials stimulated at 3 mW. **(I)** Probability of switch on trial after photostimulation by region. **(J)** Session-based performance metrics, including entropy (left), number of trials (middle) and reward rate (right). Statistical analyses were performed using two-way ANOVA for (F) and (I) and one-way ANOVA for (G) and (J), and are included in Supplementary Table.

Initially, we tested how optogenetic stimulation of noradrenergic and cholinergic axons in different brain regions may contribute to performance in matching pennies. We stimulated four regions including left and right secondary motor cortex and left and right primary visual cortex (left M2, right M2, left V1, and right V1; 40% chance of photostimulation on a given trial, equally allocated to each region) in a single session, while fixing the power at one of three levels for the session (0, 1.5, and 3 mW) **(Fig. 6E).** The most obvious effect of photostimulation was to alter the probability of a choice switch on the subsequent trial (i.e., if the mouse chose left and received a photostimulation, then the next trial it would choose right, and vice versa). With increasing power, we observed that evoking NE release in medial frontal cortex increased the tendency for the mouse to change its choice (main effect of power: F(75, 2) = 7.57, *p* = 0.001, two-way ANOVA and post-hoc Tukey test) (**Fig. 6F**; **Supplementary Table 6-1**). Curiously, this consequence of NE manipulation was equally effective for all regions stimulated (main effect of region: F(75, 4) = 0.11, *p* = 0.98; interaction of region and power: F(75, 8) = 0.05, *p* = 1.00). Similarly, there was a significant effect of photostimulation power on the switch probability (F(90, 2) = 8.39, *p* = 0.0005). We wanted to know if this photostimulation-induced propensity to alternate choices affected performance. Comparing sessions with increasing laser power, we did not detect any difference on performance metrics including entropy (NE: F(41, 2) = 0.90, *p* = 0.42; ACh: F(43, 2) = 0.20, *p* = 0.82; one-way ANOVA), number of trials performed per session (NE: F(44, 2) = 1.64, *p* = 0.21; ACh: F(43, 2) = 0.48, *p* = 0.62), or reward rate (NE: F(44, 2) = 1.09, *p* = 0.35; ACh: F(43, 2) = 0.009, *p* = 0.99) (**Fig. 6G**; **Supplementary Table 6-2**). Collectively, this photostimulation protocol increases the switch probability for both the Dbh-Cre;Ai32 and ChAT-Cre;Ai32 mice. The behavioral alterations lacked region and temporal specificity, because the choice behavior was altered on trials when any region was stimulated or even when photostimulation was absent.

We speculated that the lack of region and temporal specificity may be because the photostimulation trials were too frequent. Therefore, we modified the protocol to activate only one region at 3 mW on 10% of the trials per session **(Fig. 6H)**. With this revised photostimulation protocol, we observed a within-session difference for NE between the photostimulation and control trials (main effect of photostimulation: F(40, 1) = 5.62, *p* = 0.02, two-way ANOVA), with the mouse switching its choice significantly more on trials following optogenetic stimulation of noradrenergic axons compared to trials without (*p* = 0.022, post-hoc Tukey test; **Fig. 6I**; **Supplementary Table 6-3**). There was difference across region (main effect of region: F(40, 3) = 4.03, *p* = 0.01), with mice switching more during sessions when left M2 was stimulated compared to left and right V1 sessions (*p* = 0.06 and 0.01 respectively, post-hoc Tukey test). Although it was clear that within V1 sessions, photostimulation trials increased switching relative to control trials, suggesting that there is regional preference but not exclusivity. This is likely due to the interconnected, branching afferents of NE neurons, and our photostimulation is activating collaterals to project to multiple other cortical regions (Schwarz et al., 2015; Kim et al., 2016). With this protocol, we did not detect behavioral changes when manipulating ACh levels (main effect of region: F(48, 3) = 0.99, *p* = 0.40; main effect of stimulation: F(48, 1) = 0.07, *p* = 0.80). There was likewise no impact of the photostimulation on the whole-session performance metrics (**Fig. 6J**; **Supplementary Table 6-4**). Altogether, considering the results from both photostimulation protocols, we concluded that optogenetic stimulation of noradrenergic axons increases the switch probability on the subsequent trial, with regional preference and temporal specificity if the photostimulation was applied sparsely.

### Optogenetic elevation of frontal cortical NE in a simple choice task

Because optogenetic activation of frontal cortical noradrenergic axons promoted switching without improving reward rate, we wondered if the impact of the perturbation was specific to decision-making with competitive pressure like matching pennies or if the effect would generalize to a simplified task. We trained the same mice, after matching pennies experiments, on a simple choice task where there is no inherent benefit to switching. The structure and timing of each trial was nearly identical to matching pennies (**Fig. 7A, B**). Photostimulation (473 nm, 40 Hz, 3 mW) applied to M2 occurred on select trials starting at the choice and sustained for 1 s. Different from matching pennies, instead of a payoff matrix, reward availability followed a block structure **(Fig. 7C)**. In block 1, left choices have a 50% chance of water reward while right choices have a 50% chance of water reward paired with photostimulation. After a random number of trials, without external cue informing the mouse of the block reversal, block 2 began with the opposite action-outcome contingencies. Each mouse was tested on multiple sessions with either sham photostimulation (0 mW) or photostimulation at 3 mW in a randomized order. Example sessions illustrated the typical behavior: without photostimulation, Dbh-Cre;Ai32 and ChAT-Cre;Ai32 mice tended to stick to one option and would persist in making the same choice repeatedly (**Fig. 7D, F**). However, a Dbh-Cre;Ai32 mouse switched more frequently when the water reward was paired with photostimulation than control **(Fig. 7E)**. By contrast, a ChAt-Cre;Ai32 mouse switched rarely even when photostimulation was active **(Fig. 7G)**. Summarizing the data across all animals, mice completed a similar number of trials per session regardless of photostimulation (Dbh-Cre;Ai32: t(11) = 0.28, *p* = 0.79; ChAt-Cre;Ai32: t(8) = 0.88, *p* = 0.40; **Fig. 7H, left**). Dbh-Cre;Ai32 mice overall explored the options more by switching choices during a session with photostimulation (t(11) = -2.68, *p* = 0.02), while ChAt-Cre;Ai32 mice switched infrequently in both conditions (t(8) = 0.65, *p* = 0.53; **Fig. 7H, right**). Analyzing the data on a per-trial basis, neither strain showed a preference for the side designated for photostimulation (Dbh-Cre;Ai32: t(11) = -1.45, *p* = 0.18); ChAt-Cre;Ai32: t(8) = -1.24, *p* = 0.25, **Fig. 7I, left**). Dbh-Cre;Ai32 mice were more likely to switch on any given trial in sessions with photostimulation (t(11) = -2.55, *p* = 0.03) while ChAt-Cre;Ai32 mice showed no difference (t(8) = 0.30, *p* = 0.77; **Fig. 7I, right**). These results indicate that the evoked elevation of NE in the medial frontal cortex causes the mouse to switch choices more frequently, even though there is no preference for photostimulation per se and there is no incentive in this simple choice task for exploring.

**Figure 7.**
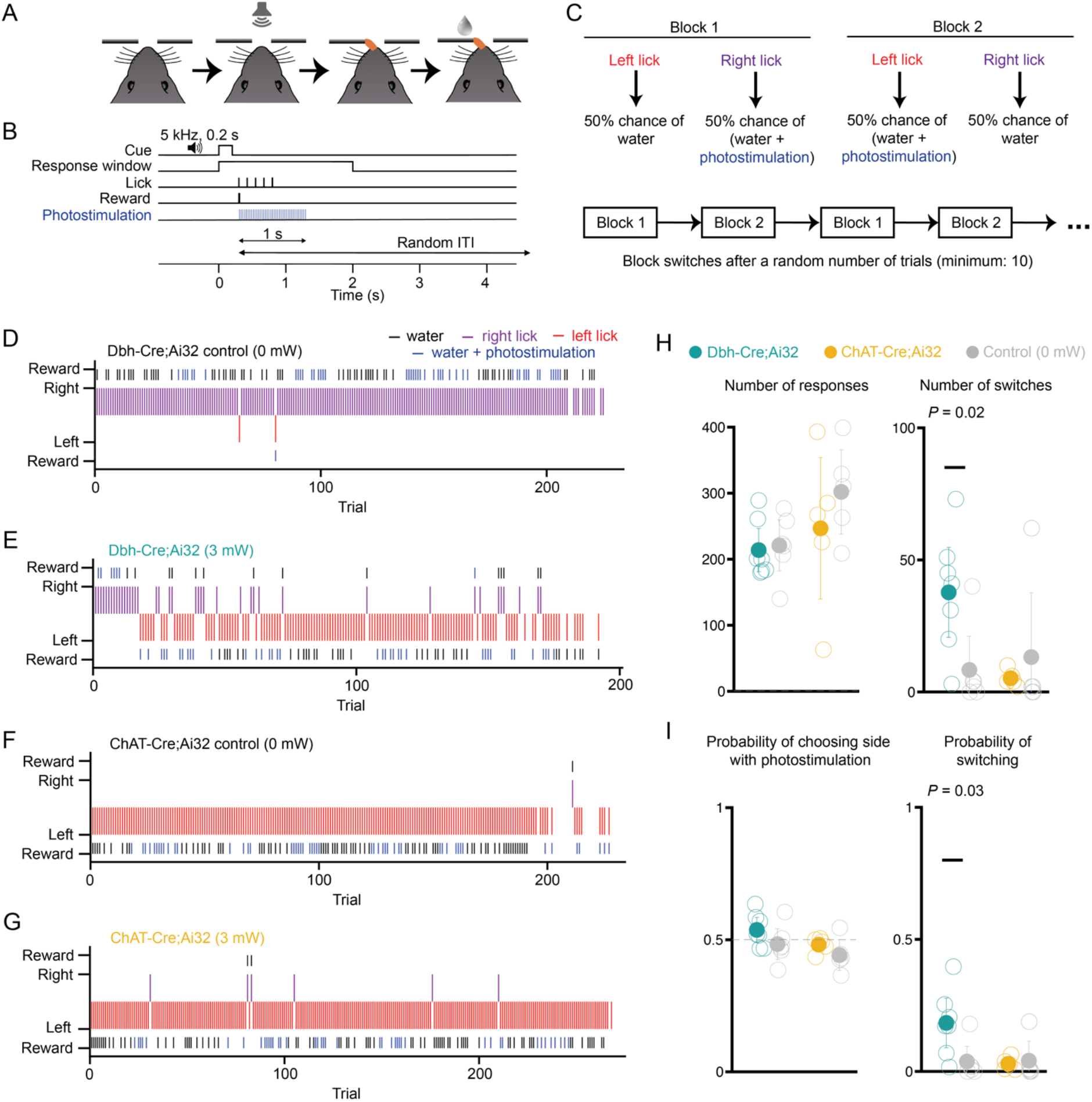
Optogenetic elevation of frontal cortical NE increases switching in a simple choice task. **(A)** Schematic of simple choice task. **(B)** Each trial, a sound cue signals the start of a response window. The first lick emitted by the animal within the window is logged as the response for that trial, and the outcome is delivered immediately according to the trial block structure. A random intertrial interval follows the outcome. On trials with photostimulation, laser turns on at time of choice and sustains until 1 s after choice. **(C)** Schematic of the trial block structure. For trials in block 1, a left lick leads to water 50% of the time and a right lick leads to water paired with photostimulation 50% of the time. For trials in block 2, action-outcome contingencies are reversed. The first block is randomly selected. The block type reverses after a random number of choices (drawn from a truncated exponential distribution, with a minimum number of 10 trials). **(D)** Example session for a Dbh-Cre;Ai32 mouse with sham stimulation. **(E)** The same mouse as in (D) but in a session with 3 mW photostimulation. **(F)** Example session for a ChAt-Cre;Ai32 mouse with sham stimulation. **(G)** Same mouse as in (F) but in a session with 3 mW stimulation. **(H)** Quantification for session-based metrics including number of responses (left) and switches (right). **(I)** Quantification for trial-based metrics including probability of choosing side designated for photostimulation (left) and probability of switching (right).

## Discussion

This study yielded three main findings. First, during a competitive game, both NE and ACh in the mouse medial frontal cortex encode task-relevant information including choice and outcome. The noradrenergic representation is more spatially segregated at the subcellular scale, whereas the cholinergic representation tends to multiplex multiple behavioral variables at the same location. Second, the decision-related NE transients are more synchronized and peak earlier than the ACh signals. Third, elevating NE levels in the medial frontal cortex promotes exploratory behavior by spurring the animal to switch choices on the subsequent trial. Together, these findings reveal distinct spatiotemporal dynamics for NE and ACh signaling in the frontal cortex, which may underpin their differential contributions to learning and decision-making.

### Imaging considerations

We can visualize the dynamic fluctuation of neuromodulator levels at subcellular resolution owing to advances in novel genetically encoded fluorescent sensors of NE and ACh. However, there are limitations to consider. Two-photon-excited fluorescence enables deep-tissue imaging, but the dense expression and relatively weak brightness of the current generation of sensors restrict the imaging depth. Therefore, we are only sampling NE and ACh transients in the supragranular layers of the medial frontal cortex. For NE, the majority of the LC inputs to the cortex resides in layer I (Swanson and Hartman, 1975). However, we are likely missing a substantial fraction of cholinergic inputs because afferents from the basal forebrain predominantly reside in infragranular layers, with 77% found in layers V and VI, and only 14% and 9% in layer I and layer II/III respectively (Henny and Jones, 2008). We obtained ∼10% change in fractional fluorescence from the most responsive ROIs and ∼1-2% change in fractional fluorescence averaged across a field of view. We were concerned that fluorescence signals may arise from motion artifact, rather than biological sources. This is why we performed the auditory evoked response and spontaneous pupillary measurements to confirm that the fluorescence signals agree with known physiological correlates of cortical NE and ACh levels.

A main finding of this study is the difference in timing, where task-related elevation of NE was significantly more aligned and peaked shortly after the decision. The 1_on_ and 1_off_ are 0.11 s and 0.58 s for GRAB_ACh3.0_ for 100 μM ACh (Jing et al., 2020), while the 1_on_ and 1_off_ are 0.09 s and 1.93 s for GRAB_NE2h_ for 100 μM NE (Feng et al., 2023). The sensors have similar rise times and GRAB_NE2h_ has slower decay time than GRAB_ACh3.0_, therefore the intrinsic kinetics of the sensors cannot account for the temporal dynamics observed in this study. There is effort to expand the color palette of the genetically encoded fluorescent sensors. Red-shifted sensors are available now for dopamine (Patriarchi et al., 2020; Zhuo et al., 2023) and has just been developed for NE (Kagiampaki et al., 2023). Future studies may leverage wavelength-shifted sensors to simultaneously monitor multiple neuromodulators at the same time, to further determine whether the spatial organization of ACh and NE transients may be coordinated and the potential interplay between different neuromodulators.

### Spatial organization of decision-related NE and ACh transients in the medial frontal cortex

Our results reveal that NE and ACh transients in the mouse medial frontal cortex occur when animals made choices and received rewards during a competitive decision-making task. Fluctuations of cholinergic and noradrenergic activities are intimately linked to pupil-associated arousal state (Reimer et al., 2016), therefore our results are consistent with prior works showing that pupil size changes are correlated with choice, outcome, and reward prediction error (Einhauser et al., 2010; de Gee et al., 2014; Van Slooten et al., 2018; Wang et al., 2022). Moreover, the finding of this study is in agreement with a recent study reporting that cholinergic basal forebrain neurons provide reinforcement signals to its axonal targets (Sturgill et al., 2020).

A notable conclusion of this study is that the decision-related signals carried by NE is more spatially distributed at the subcellular scale, whereas ACh can be modified by multiple behavioral variables at the same location. This has important implications because it suggests that ACh can influence neural plasticity and cortical computation specifically and only following more complex events that involve a conjunction of behavioral conditions. By contrast, a more segregated representation, like NE, would transmit task-related information in parallel to distinct elements of the cortical microcircuit. Representation of different behavioral variables at the subcellular scale has been observed previously in the dopamine system. Using two-photon microscopy to visualize calcium transients, individual dopaminergic axons in dorsal striatum were found to encode either locomotion onset or reward (Howe and Dombeck, 2016). The heterogeneity in behavioral correlates mapped onto genetically defined subtypes of dopaminergic neurons (Azcorra et al., 2023). Recent studies have likewise revealed subtypes of noradrenergic neurons in the LC, with distinct firing patterns during decision-making (Su and Cohen, 2022) and preferential long-range projection targets (Uematsu et al., 2017; Totah et al., 2018). There are also subtypes of cholinergic neurons in the basal forebrain, which differ in physiological properties and behavioral correlates (Laszlovszky et al., 2020). It is plausible that the spatial organization of ACh and NE transients arises due to various degree of spatial overlap of axons in the medial frontal cortex from different subtypes of NE or ACh neurons.

### Noradrenergic system promotes switching and exploratory behavior

After photostimulation of frontal cortical NE axons, animals increased tendency to switch their choice during both matching pennies and a simple choice task. Our results echo the central conclusion of an earlier study that used chemogenetics to activate LC inputs into the anterior cingulate cortex in rats, which increased behavioral variability (Tervo et al., 2014). However, unlike the earlier work, in our task there is no incentive to switch in the simple choice task, suggesting that this causally evoked behavioral change was not adaptive for improving performance. Recent study has shown that silencing the mouse anterior cingulate cortex decreases stochasticity in a foraging task (Vertechi et al., 2020), suggesting that the impact of frontal cortical NE on exploratory behavior may be bidirectional, in agreement with a theoretical proposal that NE may relate to the inverse temperature parameter in reinforcement learning (Doya, 2002).

One may ask: given the prominent task-related ACh transients, why was it that stimulating the cholinergic axons yielded no detectable change in behavior? This can be due to technical limitations, because photostimulation was applied broadly to entire brain regions. We cannot recapitulate the precise fine-scale spatiotemporal patterns observed for the neuromodulatory transients. Unlike NE, we show that ACh transients are staggered with varying peak times at different locations, which could not be mimicked by wide field optogenetic stimulation. Moreover, ACh and medial frontal cortex have roles in decision making and learning that are not captured by the behavioral tasks in this study. ACh contributes to cue-guided responses (Gritton et al., 2016) and working memory (Chudasama et al., 2004). Furthermore, medial frontal cortex including anterior cingulate cortex is involved in tracking volatility and uncertainty (Behrens et al., 2007), as well as risk aversion (van Holstein and Floresco, 2020) and belief or strategy updating (Starkweather et al., 2018; Tervo et al., 2021; Atilgan et al., 2022). These are aspects of decision-making that are not emphasized in matching pennies, which may be why optogenetic stimulation of cholinergic axons in medial frontal cortex yielded a null effect.

### Conclusion

Norepinephrine and acetylcholine are major neurotransmitters in the brain. Here, taking advantage of novel fluorescent sensors and *in vivo* two-photon microscopy, we characterized noradrenergic and cholinergic signaling in subcellular resolution in the medial frontal cortex in mice while they were engaging in a competitive decision-making game. We uncovered that decision-related events are associated with NE and ACh transients with distinct spatiotemporal dynamics. Causal manipulation of frontal cortical NE heightened exploratory behavior. Our study contributes to the emerging understanding of the functions of these neuromodulators in value-based decision-making and provides clues into why their dysfunction may underlie cognitive symptoms of neuropsychiatric disorders.

## Acknowledgements

We thank Patricia Jensen for generating and sharing the Dbh-Cre mice, Neil Savalia for help with the two-photon microscope, and Stephan Thiberge, Lucas Pinto, and David Tank for sharing the design of the laser steering system.

## Conflict of Interest

A.C.K. has been a scientific advisor or consultant for Empyrean Neuroscience, Freedom Biosciences, and Psylo. A.C.K. has received research support from Intra-Cellular Therapies. These duties had no influence on the content of this article.

## Funding source

This work was supported by NIH/NIMH R01MH112750 (A.C.K.), NIH/NIMH R21MH118596 (A.C.K.), China Scholarship Council-Yale World Scholars Fellowship (H.W.), Gruber Science Fellowship (H.K.O.), NIH training grant T32NS007224 (H.K.O.), and Cornell Engineering Learning Initiatives award (J.I.). The work used the Cornell Institute of Biotechnology’s Imaging Facility, supported by NIH 1S10RR025502 for the Zeiss LSM 710 Confocal Microscope.

## Methods

### Animal

All animal procedures were conducted in accordance with procedures approved by the Institutional Animal Care and Use Committees at Yale University and Cornell University. For imaging, adult male C57BL/6J mice were used (postnatal day 56 or older; #000664, Jackson Laboratory). For photostimulation, adult male and female Dbh-Cre;Ai32 and ChAT-Cre;Ai32 mice were used (postnatal day 42 or older). Dbh-Cre;Ai32 mice were generated by crossing B6.Cg-*Dbh^tm3.2(cre)Pjen^*/J (Tillage et al., 2020) and B6.Cg-*Gt(ROSA)26Sor^tm32(CAG-COP4*H134R/EYFP)Hze^*/J (#024109, Jackson Laboratory) (Madisen et al., 2012). ChAt-Cre;Ai32 mice were generated by crossing B6.129S-*Chat^tm1(cre)Lowl^*/MwarJ (#031661, Jackson Laboratory) (Rossi et al., 2011) and B6.Cg-*Gt(ROSA)26Sor^tm32(CAG-COP4*H134R/EYFP)Hze^*/J (#024109, Jackson Laboratory). Mice were housed in groups of three to five animals with 12/12 h light/dark cycle control (lights off at 7 P.M.).

### Surgical procedures

To prepare for imaging, animals underwent surgery for viral injection and cranial window implant. At the start of surgery, the animal was anesthetized with 2% isoflurane, which was reduced to 1-1.2% as the surgery progressed. The mouse was placed on a water-circulating heating pad (TP-700, Gaymar Stryker) in a stereotaxic frame (David Kopf Instruments). After injecting carprofen (5 mg/kg, s.c.; #024751, Butler Animal Health) and dexamethasone (3 mg/kg, s.c.; Dexaject SP, #002459, Henry Shein Animal Health), the scalp of the animal was removed to expose the skull, which was cleaned with 70% ethanol and povidone-iodine three times. For the first part of procedure, a custom-made stainless-steel head plate was glued to the skull with transparent Metabond (C&B, Parkell Inc.). For the second part of procedure, a 3-mm-diameter craniotomy was made over the longitudinal fissure (centered on AP + 1.5 mm, ML 0.0 mm relative to Bregma) using a high-speed rotatory drill (K.1070; Foredom). The dura was left intact and irrigated frequently with artificial cerebrospinal fluid (ACSF; in mM: 5 KCl, 5 HEPES, 135 NaCl, 1 MgCl2 and 1.8 CaCl2; pH 7.3) over the remainder of the procedure. The injection sites were located on the 4 vertices of a square with 0.2 mm side length, centered on a medial target within M2 (AP + 1.5 mm, ML ± 0.5 mm relative to Bregma). Either AAV9-CaMKII-GRAB_ACh3.0_ or AAV9-hSyn-GRAB_NE2h_ (titer >10^13^ GC/mL, WZ Biosciences Inc.) was infused at the four-injection site through a glass micropipette attached to a microinjection unit (Nanoject II; Drummond). Each site was injected with 46 nL of the aforementioned viruses 8 times over 2 min, at a depth of 0.4 mm from the dura. To minimize backflow of the injected solution, the micropipette was left in place for 5 min after each infusion. The cranial window consisted of one piece of 4-mm-diameter, #1 thickness prefabricated glass coverslip (#64-0720-CS-4R; Warner Instruments) and three pieces of 3-mm-diameter, #1 thickness prefabricated glass coverslips (#64-0720-CS-3R; Warner Instruments), glued together concentrically with UV-activated optical adhesive (NOA 61; Norland Products, Inc.). The window was placed on the cortical surface with the glass plug facing down with gentle downward pressure provided by a wooden stick attached to the stereotaxic frame. The window was then secured by cyanoacrylate glue and Metabond. Post-operative analgesia (carprofen, 5mg/kg, s.c.) was provided immediately and for three consecutive days following surgery. For most animals, the first and second parts of procedure were done in the same surgery, prior to behavioral training and imaging. We were concerned that this sequence prolongs the time of viral-mediated expression which may affect the signal. Therefore, for a few animals, the first and second parts of the procedure were done in separate surgeries, each with its own set of pre- and post-operative steps. The initial head plate implant allowed for training, then once the animals were proficient, we injected viruses and prepared cranial window for imaging. We did not detect differences in the two approaches and therefore present only the combined data set.

To prepare for photostimulation, the steps closely followed procedures described previously (Pinto et al., 2019). At the start of surgery, the animal was anesthetized with 2% isoflurane, which was reduced to 1-1.2% as the surgery progressed. The mouse was placed on a water-circulating heating pad (TP-700, Gaymar Stryker) in a stereotaxic frame (David Kopf Instruments). After injecting carprofen (5 mg/kg, s.c.; #024751, Butler Animal Health) and dexamethasone (3 mg/kg, s.c.; Dexaject SP, #002459, Henry Shein Animal Health), the scalp of the animal was removed to expose the skull, which was cleaned with 70% ethanol and povidone-iodine for three times. After removing the scalp, the skull was lightly polished using acrylic polish kit (S23-0735, Pearson Dental) to remove residual tissue. A custom-made stainless steel headplate (eMachineShop) was glued onto the skull with Vetbond and the center well filled with transparent Metabond (1 scoop purple powder, 7 drops base, 2 drops catalyst; C&B, Parkell) to obtain a ∼1 to 2 mm thick layer. After waiting for about 20 minutes for the Metabond to cure, the surface of the Metabond layer was polished with progressively finer bits from the acrylic polish kit. After polishing, the well was covered with a very thin layer of clear nail polish (72180, Electron Microscopy Services) and allowed to dry fully. Post-operative analgesia (carprofen, 5mg/kg, s.c.) was provided immediately and for three consecutive days following surgery. Animals were implanted with this clear skull cap for at least 2 weeks before the start of behavioral training.

### Behavioral setup

The same training apparatus was used in our prior studies (Siniscalchi et al., 2019; Wang et al., 2022). Detailed instruction to construct the apparatus is available at https://github.com/Kwan-Lab/behavioral-rigs. Briefly, the mouse with a head plate implant was head-fixed to a stainless-steel holder (eMachineShop). The animal, restrained by an acrylic tube (8486K433; McMaster-Carr), was able to adjust its posture with limited gross movements. Two lick spouts made of blunted 20-gauge stainless-steel needles were positioned in front of the subject near its mouth. The animal indicated its choice by licking the spout with its tongue. The contact with the lick spout formed a closed loop with wires that were soldered onto the spout and a battery-powered lick detection electronic circuit, which generated an output electrical signal. A computer received the signal via a data acquisition unit (USB-201, Measurement Computing) and logged it with the Presentation software (Neurobehavioral Systems). Two solenoid fluid valves (MB202-V-A-3–0-L-204; Gems Sensors & Controls) controlled the water delivery from the two lick ports independently. The amount of water was calibrated to ∼4 μl per delivery by adjusting the duration of the electrical pulse sent by the Presentation software through a second data acquisition unit (USB-201, Measurement Computing). The sound cue signaling the trial start was played by two speakers (S120, Logitech) placed in front of the mouse. The whole setup was placed inside an audiovisual cart with walls covered by soundproof acoustic foams (5692T49, McMaster-Carr).

### Two-photon imaging

The behavioral setup described above was placed under the two-photon microscope. The two-photon microscope (Movable Objective Microscope, Sutter Instrument) was controlled using ScanImage software 5.1. The excitation source was a Ti:Sapphire femtosecond laser (Chameleon Ultra II, Coherent). Laser intensity was controlled by a Pockels cell (350-80-LA-02, Conoptics) and an optical shatter (LS6ZM2; Uniblitz/Vincent Associates). The beam was focused onto the sample with a 20×, N.A. 1.00 water immersion objective (N20X-PFH, Thorlabs via Olympus). The time-averaged excitation laser intensity was 120–180 mW after the objective. To image fluorescence transients from GRAB_NE2h_ or GRAB_ACh3.0_ sensors, excitation wavelength was set at 920 nm and emission was collected from 475–550 nm with a GaAsP photomultiplier tube. Time-lapse images were acquired at a resolution of 256 × 256 pixels and a frame rate of 30.03 Hz using bidirectional scanning with resonant scanners. To synchronize behavioral and imaging data, a TTL pulse was sent by the Presentation software at the beginning of each trial from the USB-201 board of the behavioral system that controlled the water valves. The imaging system used the TTL pulse as an external trigger to initiate the imaging acquisition.

### Photostimulation

The photostimulation apparatus had a design based on earlier work (Pinto et al., 2019) and is the exact same configuration used in previous study (Atilgan et al., 2022). Briefly, a 473nm fiber-coupled laser (473 nm, 75mW; Obis LX, Coherent) was controlled by a pulse sequence generator (Pulse Pal, Sanworks). The fiber output was directed to a galvanometer-galvanometer scanner (6210H, Cambridge Technologies), which were driven by power supplies (SPD-3606, Cole-Parmer) and installed in a 60-mm cage system (ThorLabs). The excitation beam then passes through an F-theta scan lens (f = 160 mm; FTH160-1064-M39, ThorLabs) and is directed onto the animal’s head. Calibration of the laser beam’s position relative to bregma is achieved by visualizing the cortical surface using a monochromatic camera (Grasshopper3; GS3-U3-23S6M-C, Point Grey) with a telecentric lens (TEC-55, Computar). A blue LED (470 nm) aimed at the animal’s head was used as a masking light. Control of the laser, scanner, camera, and LED was executed through a data acquisition board (PCIe-6343, National Instruments) utilizing custom software written in MATLAB (Mathworks). The behavior setup described above was placed under the photostimulation apparatus. The entire system is housed inside a custom T-slot frame box (80/22 LLC), shielded with soundproof foam panels, on a vibration isolation table (CleanTop 781-651-02R, TMC).

### Matching pennies

Animals were trained to play the matching pennies game with a component opponent (Wang et al., 2022). All procedures were written using the programming language in the Presentation software. The animals were fluid restricted with water provided during the daily behavioral session. On the days when the subjects were not trained (typically 1 day per week), a water bottle was placed in the home cage, allowing for ad libitum water access for 5 minutes.

Animals were trained in 3 phases. For phase 1 (2 days), the animals were habituated to the behavior apparatus. They may lick either spout for water. A water reward would be delivered after every lick at the corresponding spout with a minimal time interval of 1 s. The session would terminate after the animal collected 100 rewards. For phase 2 (approximately 4 weeks), the animals were trained to follow the trial structure and withhold impulsive licks before the trial started. In each trial, a 5-kHz sound cue lasting for 0.2 s signaled the start of the trial. Then the animal was given a 2-s window to lick either port. The 2-s response window would give a naïve mouse more time to act when they had not learnt the trial timing, therefore helping the animals to acquire the task faster. Once the first lick was detected, the 2-s response window would be terminated immediately. A water reward would be presented at the corresponding spout, following which a fixed 3-s period was presented for the animal to collect the reward. In the trials when the animal did not lick, the 3-s interval was still presented in full. A random intertrial interval (ITI) began after the 3-s consumption window. A number was drawn from a truncated exponential distribution with lambda=0.333 and boundaries of 1 and 5, which was used as the duration of the ITI in seconds. If one or more licks were detected during the ITI, an additional ITI with duration redrawn from the same distribution would be appended to the end of current ITI, with a maximum of 5 ITIs. After the ITIs ended, the next trial would begin. The animal would be advanced into phase 3 to play the matching pennies game when the average number of ITI draws per trial was lower than 1.2 for 3 consecutive sessions. In phase 3 (approximately 4 weeks), the animals were trained to play the matching pennies game against a computer opponent whose behavior was controlled by a script written in the programming language of the Presentation software. The trial timing is the same as phase 2: each trial begins with a 5-kHz, 0.2-s sound cue. Within a 2-s response window, the animal indicated its choice by licking either the left or right spout. A water reward would be delivered in the corresponding spout if the animal chose the same choice as the computer. Otherwise, there would be no reward. The computer opponent was programmed to provide competitive pressure in a way the same as “algorithm 2” described in previous studies (Barraclough et al., 2004; Lee et al., 2004; Wang et al., 2022). Specifically, the computer opponent kept a record of all the animal’s past choices and outcomes within the current session and ran 9 binomial tests on the conditional probability of the animal choosing left given the sequence of previous N choices (N=0-4) and previous M choices and outcomes (M = 1-4), against the null hypotheses that the conditional probabilities of the animal choosing left was 0.5. If at least one of the tests rejected the null hypotheses with alpha < 0.05, the computer then chose right with the significant conditional probability that was most biased from 0.5. If none of the null hypothesis was rejected, the computer randomly generated either choice with equal probabilities. The animal could play for as many trials as it desires, and a session would terminate when no response was detected for 10 consecutive trials. Mice reached stable performance when they played matching pennies for 3 consecutive sessions with a minimum of 40% reward rate.

Initially, mice were trained in dedicated behavioral setups. After reaching criterion, animals were trained to play the same matching pennies game in the behavioral setup within the two-photon imaging or photostimulation rig. They would be deemed to have adapted when mice played matching pennies for 3 consecutive sessions with a minimum of 40% reward rate, which was when imaging or photostimulation experiments would commence.

### Matching pennies and photostimulation

During matching pennies, laser was turned on for photostimulation (frequency: 40 Hz; pulse duration: 0.1 ms) on select trials from onset of cue to 1 s after choice was made (i.e., first lick within response window). Photostimulation was applied to one of four possible locations: left secondary motor cortex (M2-L; +1.5 mm AP, -0.3 mm ML from bregma), right secondary motor cortex (M2-R; +1.5, +0.3), left primary visual cortex (V1-L; -3.0, -2.0), or right primary visual cortex (V1-R; -3.0, +2.0). On trials without photostimulation, the masking blue LED would be turned on from onset of cue to 1 s after choice was made. For the “varying power” paradigm, photostimulation occurred on 40% of the trials randomly with 10% allocated to each region (M2-L, M2-R, V1-L, and V1-R). One power was used for a session, but power changed across sessions in a pseudorandom order between 0, 1.5, and 3 mW. For the “varying region” paradigm, photostimulation occurred on 10% of the trials at 3 mW. One region was tested for a session, but region changed across sessions in a pseudorandom order between M2-L, M2-R, V1-L, and V1-R.

### Simple choice task

Each trial begins with a 5-kHz, 0.2-s sound cue. Within a 2-s response window, the animal indicated its choice by licking either the left or right spout. In trial type 1, left spout has 50% chance of delivering water and photostimulation (from choice to 1 s after choice was made) while right spout has 50% chance of delivering only water. In trial type 2, left spout has 50% chance of delivering only water while right spout has 50% chance of delivering photostimulation and water. A session begins with a first trial of trial type 1 or 2 randomly. The next trial has a 1/11 probability to switch trial type. There were no external stimuli beyond the probabilistic photostimulation and water delivery to inform the mouse of the trial type switch. The animal could play for as many trials as it desires, and a session would terminate when no response was detected for 20 consecutive trials.

### Pupillometry

A monochrome camera (GigE G3-GM11-M1920, Dalsa) with a 55 mm telecentric lens (TEC-55, Computar) was aimed at the eye of the animal contralateral to the hemisphere where imaging was performed. Video was acquired at 20 Hz. The computer running the Presentation software sent TTL pulses every 30 s to another computer controlling the camera through a USB data acquisition device (USB-201; Measurement Computing). The timestamp of the TTL pulse was logged by MATLAB 2019b (MathWorks) with a custom script, such that the video could be aligned to behavioral events post hoc. The computer running the Presentation software sent TTL pulses every 30 s to the two-photon microscope to trigger imaging. Each session lasted 30 minutes. Animals were tested either naïve or after going through the entire behavioral training protocol for matching pennies.

### Auditory evoked responses

This measurement relied on the same behavioral setup as the matching pennies. A 4-kHz, 50-ms auditory stimulus was played at the beginning of each trial. A random ITI was presented following the stimulus. The duration of the ITI in seconds was drawn from a continuous uniform distribution with boundaries of 1 and 4. The next trial would begin after the ITI. Each session lasted 200 trials.

### Histology

Following imaging experiments, mice were transcardially perfused with chilled phosphate-buffered saline (PBS) followed by formaldehyde solution (4% in PBS). The brains were then fixed in 4% formaldehyde solution for 1 hour before they were transferred to 30% sucrose solution at 4*°*C. After about 24 hours, the brains were cut into 50-μm thick coronal sections with a vibratome (VT 1000S, Leica). The brain sections were washed 3 times with PBS solution before being immersed with chicken anti-GFP antibody (1:500; ab13970, Abcam) for 12 hours at 4*°*C. Then Alexa 488-conjugated goat anti-chicken secondary antibody (1:50; ab209487, Abcam) was used to label the primary antibody for 3 hours at room temperature. The sections were then mounted with DPX and imaged with an inverted wide-field fluorescence microscope. Following photostimulation experiments, mice were transcardially perfused with chilled PBS then formaldehyde solution (4% in PBS). The brains were fixed in 4% formaldehyde solution for 24 hours at 4*°*C before being transferred to PBS. Then brains were processed using the vibratome into 30-μm thick coronal sections. Sections were washed 5 times with PBS and incubated in PBS with 0.3% Triton X (PBST) for 20 minutes at room temperature. Slices were blocked with 10% normal goat serum in PBST for 1 hour at room temperature followed by incubation with primary anti-GFP antibody (1:200, ab290, Abcam) in 10% normal goat serum in PBST at 4*°*C overnight. Sections were washed 3 times in PBS then incubated with Alexa 488-conjugated goat anti-rabbit secondary antibody (1:500; ab150077, Abcam) for 2 hours at room temperature. Slices were mounted with Vectashield Antifade Mounting Medium with DAPI (H-1200-10, VectorLabs) and imaged with a Zeiss LSM 710 confocal microscope.

### Preprocessing of matching pennies data

To quantify the randomness in the animals’ choices, the 3-choice entropy of the choice sequence is calculated by:

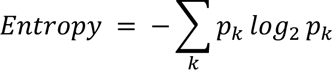

where *p_k_* is the frequency of occurrence of a 3-choice pattern in a session. Because there were 2 options to choose from, there were 2^3^ = 8 potential patterns possible (e.g., left-left-left, left-left-right, left-right-left, etc.). The maximum value for entropy is 3 bits. For matching pennies, the animals tended to select the same option for around 30 trials towards the end of each session. The 3-choice entropy over a moving 30-trial window was calculated for each session, and the MATLAB function *ischange* was used to fit with a piecewise linear function. The trials after the fitted curve fell below a value of 1 were discarded to exclude the repetitive trials in the analyses. In cases where the curve recovered to a value greater than 1 after it dropped below 1, or if it never fell below a value of 1, the entire session was used for analysis.

### Preprocessing of pupillometry data

The preprocessing of pupillometry data was similar to a previous work (Wang et al., 2022). To extract the diameters of the pupil from the video recordings, we used DeepLabCut (DLC) 2.0 (Mathis et al., 2018; Nath et al., 2019). Five labels including the central, uppermost, leftmost, lowermost, and rightmost points of the pupil were manually labelled by the experimenter on a small subset of the video frames. The annotated frames were used to train DLC to automatically label the 5 points on the remainder of the video. The absolute pupil diameter was calculated by taking the distance between the leftmost and rightmost labels. The other labels were not considered because we found the labelling of the lowermost points were interfered by the lower eyelid, resulting in an inaccurate estimation. The absolute pupil diameter was passed through a 4 Hz lowpass filter with the MATLAB function *lowpass*. Using the MATLAB function *isoutlier*, we detected and deleted any data points that were greater than 3 scaled median absolute deviation (MAD) from the median. The baseline of the signal was computed with a 10-minute moving window, which was used to convert the signal to z-score to account for drift over a session.

### Preprocessing of imaging data

Time-lapse images were processed for x-y motion correction using customized MATLAB scripts based on NoRMCorre (Pnevmatikakis and Giovannucci, 2017). The field of view (FOV) spans 142.66×142.66 μm, or 256×256 in pixels. For analysis, we only use the 124.83×124.83 μm, or 224×224 pixels, portion at the center of the FOV to avoid artifacts near the FOV edges due to motion correction processing. The analyzed region was divided into 28×28 grids. Each grid was a 4.46×4.46 μm, or 8×8 pixels, square, which was considered as a region of interest (ROI). We decided to use grids to subdivide the FOV as an unbiased way to analyze spatial dependence, rather than neural morphology because the GRAB_NE2h_ and GRAB_ACh3.0_ sensors labeled primarily neuropils that are indistinct for manual drawing of ROIs. For each ROI, the fractional change in fluorescence, *ΔF/F(t)*, was calculated as:

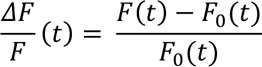

where *F*_0_(*t*) is the baseline fluorescence as a function of time, which was the 10^th^ percentile of the fluorescent intensity within a 2-minute running window centered at time *t*.

### Analysis of imaging data – Peristimulus time histogram

To obtain trial-averaged activity traces (peristimulus time histogram, PSTH) aligned to the cue onset for different trial types, we first aligned *ΔF/F(t)* traces based on their timing relative to the cue onset of the corresponding trials and then took the mean across traces. We then estimated the average and 95% confidence interval over mean traces of all recorded ROIs with bootstrapping procedure to get the average PSTH for a single session.

### Analysis of imaging data – Linear regression

To determine how fluorescence signals may relate to task-related variables such as choices and outcomes, we used a multiple linear regression equation adapted from our previous study (Wang et al., 2022):

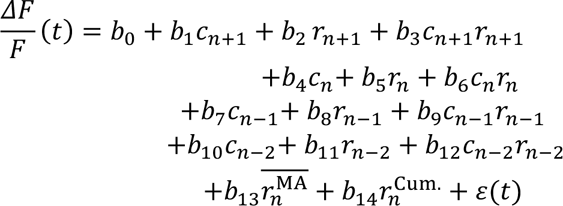

where 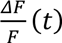 is the fractional changes in fluorescence at time *t* in trial *n*, *C*_*n*>+1_, *C_n_*, *C*_*n*–1_, *C*_*n*–2_ are the choices made on the next trial, the current trial, the previous trial, and the trial before the previous trial, respectively, *r_n_*, *r*_*n*–1_, *r*_*n*–2_ are the outcomes for the next trial, the current trial, the previous trial, and the trial before the previous trial, respectively, *b*_0_,…, *b*_14_) are the regression coefficients, and ε(*t*) is the error term. Choices were dummy-coded as 0 for ipsilateral responses and 1 for contralateral responses. Outcomes were dummy-coded as 0 for no-reward and 1 for reward. For the last 2 predictors, 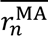 is the average reward over the previous 20 trials, given by the equation:

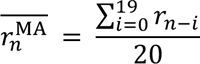

And 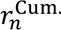 is the normalized cumulative reward during the session, calculated by:

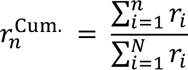

where *n* denotes the current trial number and *N* is the total number of trials in the session.

For each session, the regression coefficients were determined by fitting the equations to data using the MATLAB function *fitlm*. The fit was done in 100-ms time bins that span from -3 to 5 s relative to cue onset, using mean *ΔF/F* within the time bins. For a given predictor and an ROI, if the regression coefficients were significant (*P* < 0.01) for at least 3 consecutive or 10 total time points, the ROI was considered significantly modulated by the predictor. To summarize the results, for each predictor, we calculated the proportion of ROIs in which the regression coefficient was significantly different from zero (*P* < 0.01). To determine if the proportion was significantly different from chance, we performed a chi-square test against the null hypothesis that there was a 1% probability that a given predictor was mischaracterized as significant by chance in a single session.

### Analysis of imaging data – Hierarchical clustering

To analyze the degree of similarity in task-related activity across ROIs, within each session, we applied hierarchical clustering on the regression coefficients for the current outcome for the ROIs that were significantly modulated by the current outcome. We clustered the ROIs within each session into 2 clusters based on the cross-correlation of the regression coefficients within - 3 to 5 seconds from cue using the MATLAB function *clusterdata*. The optimal number of clusters was validated with silhouette analysis. Note that the *clusterdata* function would always cluster any given dataset into two clusters, yet in some sessions the data were better fitted with only one cluster. Therefore, we identified two distinct groups *post hoc* based on the direction of modulation, then placed every cluster identified by *clusterdata* into either of these two groups as follows: first we calculated the mean coefficients as a function of time of each cluster over different ROIs; we then calculated the area under the curve of the average coefficients between 0-2 s from cue. Clusters with a positive area under the curve were considered Group 1, and clusters with a negative area under the curve were considered Group 2. Once all the ROIs from all sessions were categorized into either group, we then identified the ROIs that were significantly modulated by current choice and/or current reinforcer (choice-outcome interaction) within the groups. For visualization, the ROIs were sorted by the center-of-mass of the regression coefficients for the current outcome. The hierarchical clustering procedures described above were performed for GRAB_NE2h_ and GRAB_ACh3.0_ data separately.

### Analysis of imaging data – Temporal dynamics of the task-related activity

To quantify the temporal dynamics of the task-related activity, the time-to-peak value was calculated for each ROI as the duration from cue onset time to the peak coefficient time (the time when the regression coefficient reached the maximum magnitude). Afterwards, the median and variance of time-to-peak and the median of peak-value were taken for each session. This quantification was performed for different regression coefficients (e.g., choice, outcome, and reinforcer) separately.

### Analysis of imaging data – Information encoding of single ROIs

To examine the spatial patterns of task-related activity, we calculated the average regression coefficients over time for the current choice, current outcome, and current reinforcer (choice-outcome interaction) separately for each session. We used the Pearson’s chi-square test for independence to test whether the event that a given ROI was modulated by one predictor was independent from the event that the same ROI was modulated by another predictor.

To compare the level of information integration between GRAB_NE2h_ and GRAB_ACh3.0_ data, we calculated 6 conditional probabilities that an ROI is modulated by one variable (*v_1_*) given that the same ROI is modulated by another variable *v_2_* for each session, given by the equation:

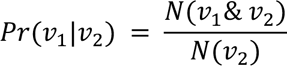

where *N*(*v*_1_& *v*_2_) denotes the number of ROIs that were modulated by *v*_1_ and *v*_2_; *N*(*v*_2_) denotes the number of ROIs that were modulated by *v*_1_ and *v*_2_ can be either *C_n_*, *r_n_*, or *x_n_* (*v*_1_≠ *v*_2_). The median test was used to determine if the conditional probabilities of an ROI was modulated by one predictor given the ROI was modulated by another predictor were significantly different between the GRAB_NE2h_ and GRAB_ACh3.0_ data, because the median test is more robust with small sample size and less sensitive to asymmetry.

### Analysis for auditory evoked responses

To obtain the mean auditory evoked responses, we first aligned the *ΔF/F(t)* based on the times of auditory stimulus onset, then calculated the peri-stimulus time histogram (PSTH) over all 200 trials for each ROI. To examine the temporal dynamics of the auditory evoked response, the time-to-peak-*ΔF/F* value was calculated for each ROI as the duration from cue onset time to the peak-*ΔF/F* time (the time when the PSTH reached the highest value). The decay time was the decay constant obtained by fitting the PSTH from the peak time to the end of trial (3 s after the cue time) to single exponential decay. Afterwards, the median of time-to-peak and decay time were taken for each session.

### Analysis for pupillary fluctuation

For plotting example traces of fluorescent and pupil signals, we smoothed the *ΔF/F(t)* and pupil z-score traces using a Gaussian kernel with the MATLAB function *smooth*. To characterize the relationship between *ΔF/F(t)* and pupil z-score, we first determine the cross-correlation between *ΔF/F(t)* and pupil z-score from lag of -2 to 2 seconds for each ROI, then determine the maximum cross-correlation value within this lag range, and finally tabulate a histogram for the maximum coefficients for all ROIs separately for GRAB_NE2h_ and GRAB_ACh3.0_ data.

### Analysis for photostimulation

For the mouse’s probability to stay on the next trial, a multi-variate ANOVA was performed for each condition (ChAt-Cre;Ai32 or Dbh-Cre;Ai32) to determine the effect of region and power or stimulation. Post-hoc Tukey test was used as needed. For analyzing session-wide metrics such as entropy, number of trials completed, and reward rate, a one-way ANOVA was performed for each condition to determine the effect of power or region. Post-hoc Tukey test was used as needed. For the simple choice task, a t-test was performed to compare sessions with and without stimulation (3 mW vs. 0 mW) for each condition on metrics of number of responses/switches, probability of choosing side with stimulation, and probability of switching on the next trial.

### Animal numbers

For two-photon imaging, the data set for matching pennies included 47 sessions from 5 animals expressing GRAB_ACh3.0_ and 38 sessions from 4 animals expressing GRAB_NE2h_. Auditory evoked responses included 7 sessions from 5 animals expressing GRAB_ACh3.0_ and 7 sessions from 7 animals expressing GRAB_NE2h_. Spontaneous pupil fluctuation included 10 sessions from 5 animals expressing GRAB_ACh3.0_ and 7 sessions from 7 animals expressing GRAB_NE2h_. For photostimulation, the data set came from 7 ChAt-Cre;Ai32 and 6 Dbh-Cre;Ai32 mice. The varying power paradigm included 91 sessions from 7 ChAt-Cre;Ai32 mice and 76 sessions from 6 Dbh-Cre;Ai32 mice. The varying region paradigm included 49 sessions from the same 7 ChAt-Cre;Ai32 mice and 41 sessions from the same 6 Dbh-Cre;Ai32 mice. The simple choice task included 8 sessions from 4 ChAt-Cre;Ai32 mice and 12 sessions from 6 Dbh-Cre;Ai32.

### Code accessibility

The data and code that support the findings of this study will be made publicly available at https://github.com/Kwan-Lab.

## Supplementary Material

**Supplementary Table 4-1.** Statistical results for the level of information integration in GRAB_NE2h_ and GRAB_ACh3.0_ data.

|  | **NE**<br>**(Number of**<br>**ROIs, %)** | ***p*-value**<br>**(Chi-square**<br>**test)** | **ACh**<br>**(Number of**<br>**ROIs, %)** | ***p*-value**<br>**(Chi-square**<br>**test)** |
| --- | --- | --- | --- | --- |
| **Choice-outcome**<br>**(Chance level)** | 2,917, 10.0%<br>(9.0%) | 0 | 8,357, 22.7%<br>(16.3%) | 0 |
| **Outcome-reinforcer**<br>**(Chance level)** | 2,819, 9.5%<br>(8.4%) | 0 | 3,316, 9.0%<br>(6.0%) | 0 |
| **Choice-reinforcer**<br>**(Chance level)** | 2,153, 7.3%<br>(3.5%) | 0 | 2,252, 6.1%<br>(2.7%) | 0 |

**Supplementary Table 4-2.**
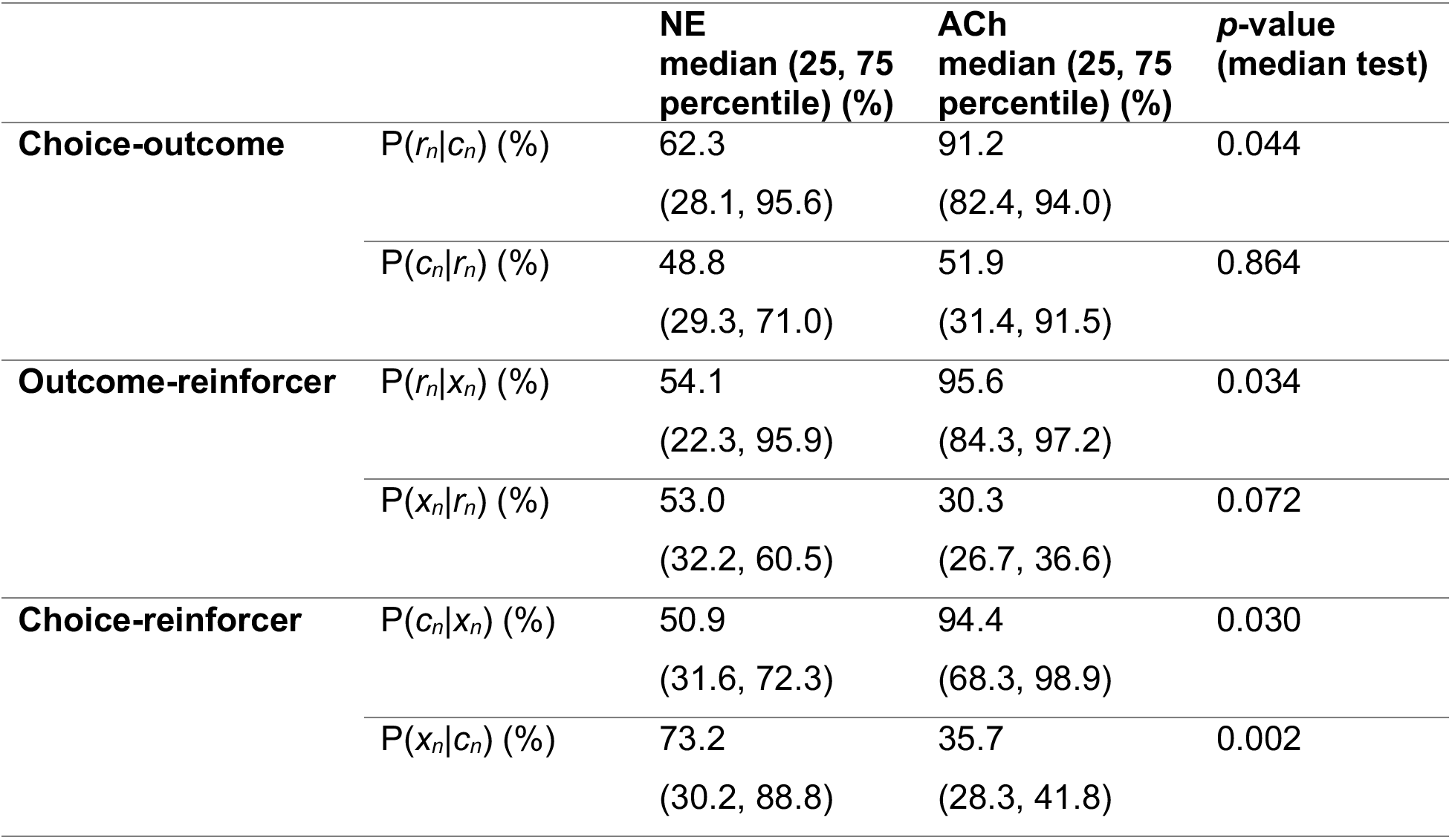
Statistical results for the conditional probabilities P(*v_1_*|*v_2_*)

**Supplementary Table 5-1.** Statistical results for the temporal parameters of different groups.

|  |  |  | **NE**<br>**median (25,**<br>**75 percentile)** | **ACh**<br>**median (25,**<br>**75 percentile)** | **p-value**<br>**(rank sum**<br>**test)** |
| --- | --- | --- | --- | --- | --- |
| **Outcome** | Group<br>1 | Median time-to-peak (s) | 0.80<br>(0.75, 0.85) | 1.95<br>(1.35, 2.45) | $1.54 \times 10^{-13}$ |
|  |  | Variance time-to-peak (s<sup>2</sup>) | 0.21<br>(0.08, 1.04) | 0.82<br>(0.28, 1.37) | 0.002 |
|  |  | Median peak-value (a.u.) | 0.07<br>(0.05, 0.09) | 0.05<br>(0.04, 0.06) | 0.003 |
| | Group<br>2 | Median time-to-peak (s) | 0.75<br>(0.75, 0.85) | 0.95<br>(0.85, 1.15) | $1.43 \times 10^{-4}$ |
|  |  | Variance time-to-peak (s<sup>2</sup>) | 0.12<br>(0.01, 0.60) | 0.37<br>(0.12, 0.58) | 0.108 |
|  |  | Median peak-value (a.u.) | -0.04<br>(-0.06, 0.03) | -0.03<br>(-0.04, -0.02) | 0.002 |
| **Choice** | Group<br>1 | Median time-to-peak (s) | 0.55<br>(0.35, 0.80) | 1.85<br>(1.25, 2.95) | $1.06 \times 10^{-7}$ |
|  |  | Variance time-to-peak (s<sup>2</sup>) | 0.62<br>(0.02, 1.46) | 1.50<br>(0.33, 3.15) | 0.003 |
| | | Median peak-value (a.u.) | -0.04<br>(-0.07, -0.02) | 0.03<br>(0.01, 0.04) | $3.53 \times 10^{-14}$ |
|  | Group<br>2 | Median time-to-peak (s) | 0.55<br>(0.38, 0.65) | 1.55<br>(0.80, 2.08) | 0.022 |
|  |  | Variance time-to-peak (s<sup>2</sup>) | 0.01<br>(0.00, 0.15) | 0.58<br>(0.21, 1.32) | 0.001 |
| | | Median time-to-peak (a.u.) | 0.02<br>(0.00, 0.05) | -0.03<br>(-0.04, -0.02) | $1.88 \times 10^{-7}$ |

**Supplementary Table 6-1.**
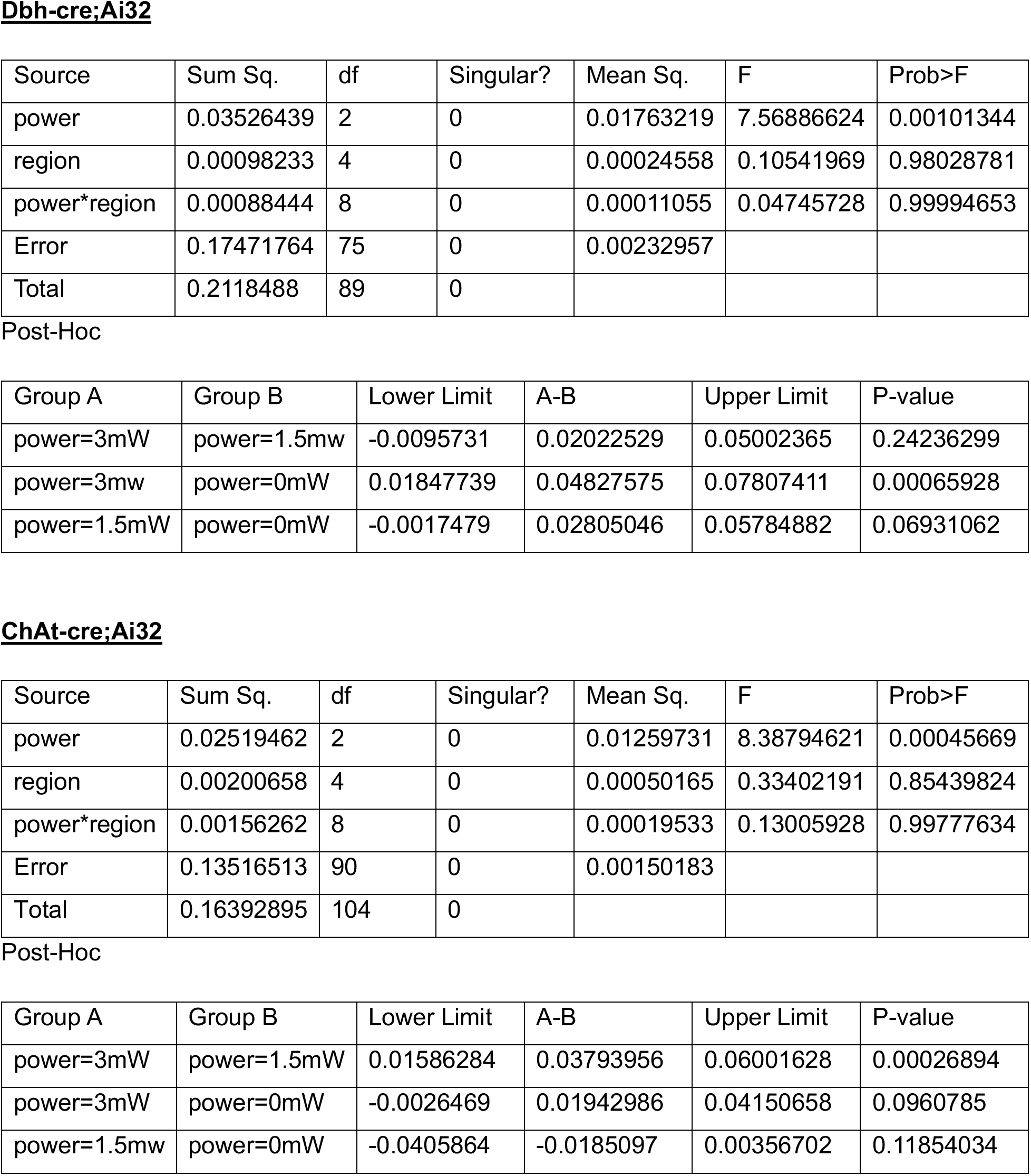
Statistical results for the effect of region and power on probability to switch, related to Fig. 6F.

**Supplementary Table 6-2.**
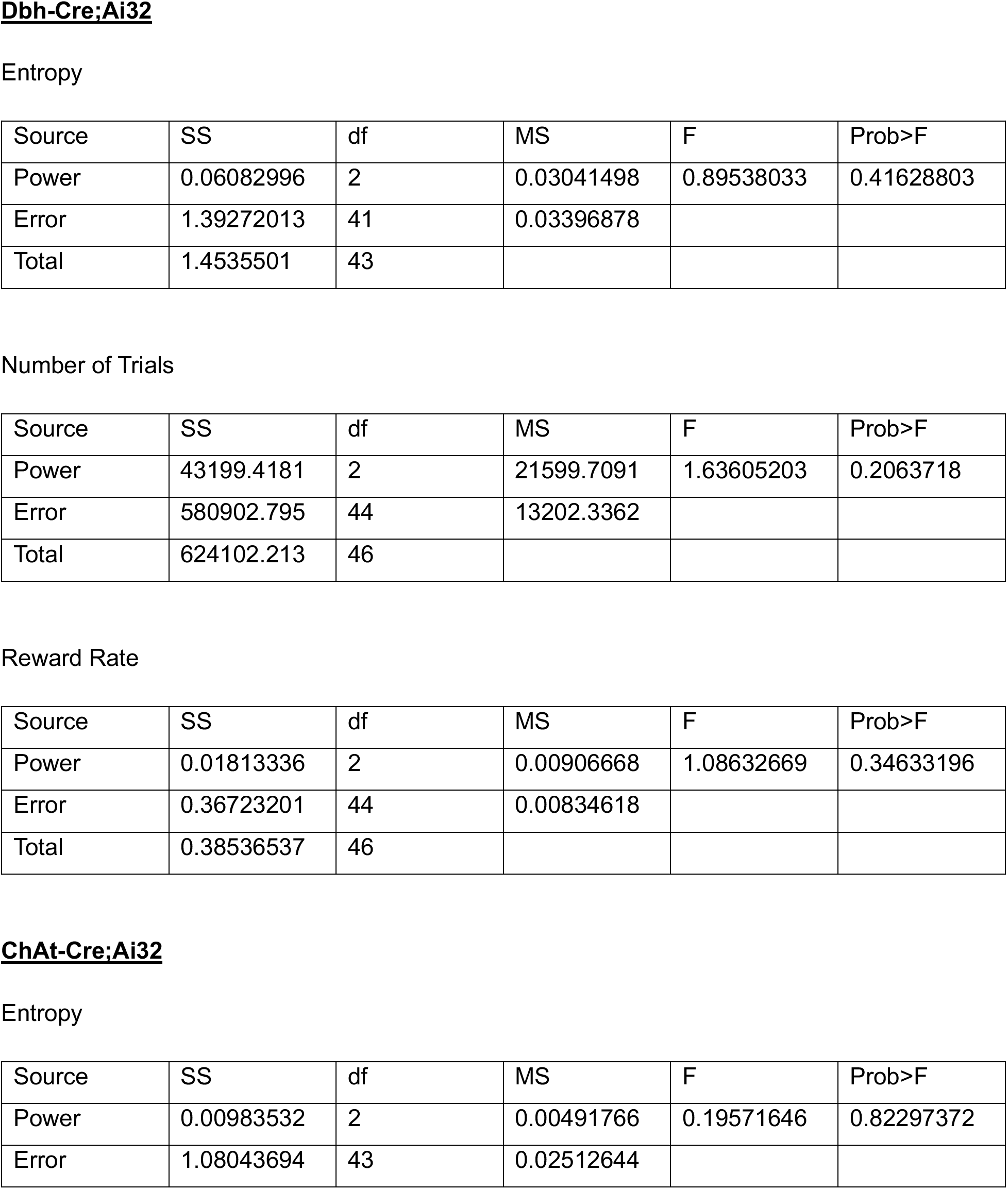
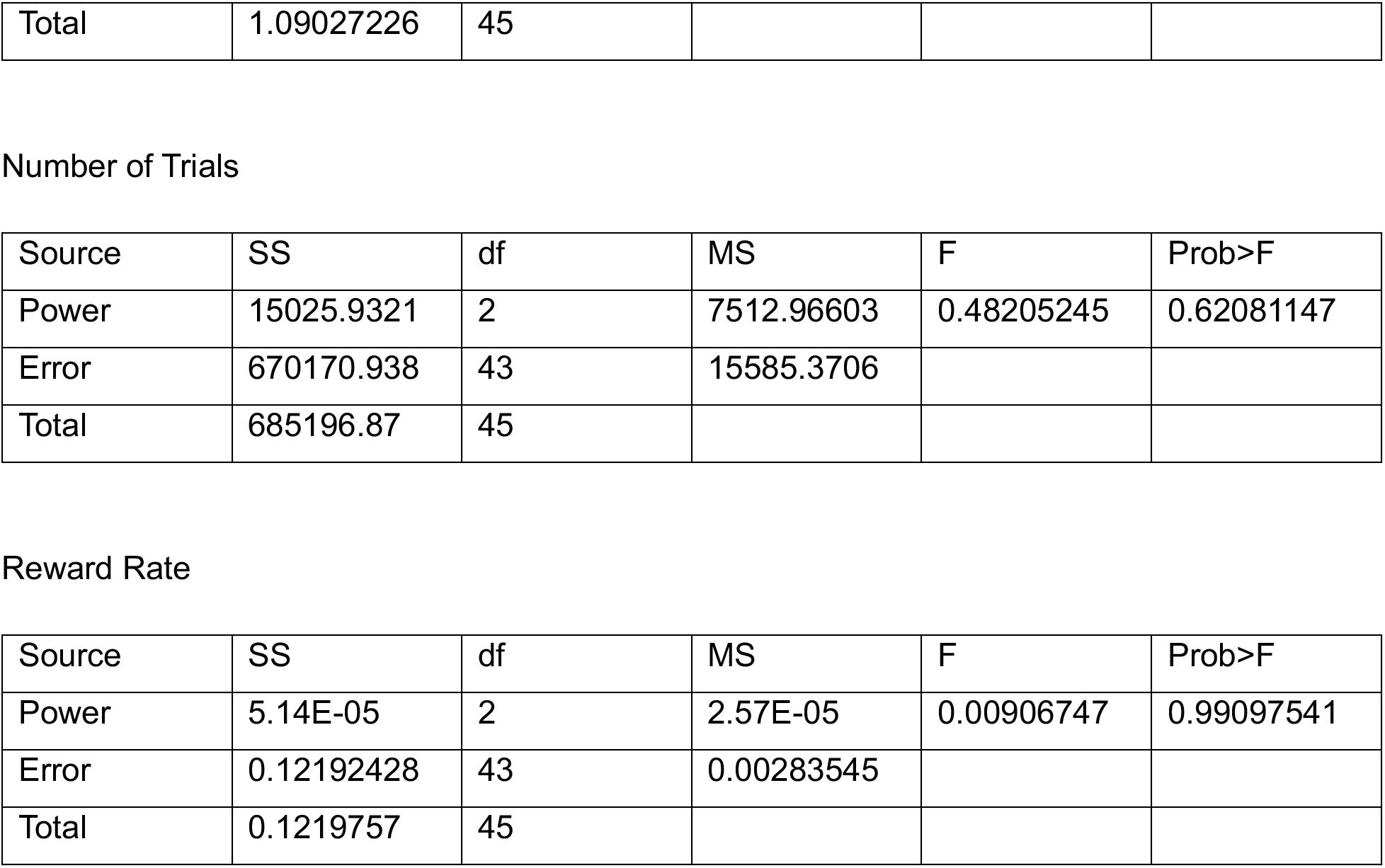
Statistical results for the effect of power on overall session, related to Fig. 6G.

| Source | SS | df | MS | F | Prob>F |
| --- | --- | --- | --- | --- | --- |
| Power | 0.06082996 | 2 | 0.03041498 | 0.89538033 | 0.41628803 |
| Error | 1.39272013 | 41 | 0.03396878 |  |  |
| Total | 1.4535501 | 43 |  |  |  |

Number of Trials
| Source | SS | df | MS | F | Prob>F |
| --- | --- | --- | --- | --- | --- |
| Power | 43199.4181 | 2 | 21599.7091 | 1.63605203 | 0.2063718 |
| Error | 580902.795 | 44 | 13202.3362 |  |  |
| Total | 624102.213 | 46 |  |  |  |

Reward Rate
| Source | SS | df | MS | F | Prob>F |
| --- | --- | --- | --- | --- | --- |
| Power | 0.01813336 | 2 | 0.00906668 | 1.08632669 | 0.34633196 |
| Error | 0.36723201 | 44 | 0.00834618 |  |  |
| Total | 0.38536537 | 46 |  |  |  |

**ChAt-Cre;Ai32** Entropy
| Source | SS | df | MS | F | Prob>F |
| --- | --- | --- | --- | --- | --- |
| Power | 0.00983532 | 2 | 0.00491766 | 0.19571646 | 0.82297372 |
| Error | 1.08043694 | 43 | 0.02512644 |  |  |
| Total | 1.09027226 | 45 |  |  |  |

Number of Trials
| Source | SS | df | MS | F | Prob>F |
| --- | --- | --- | --- | --- | --- |
| Power | 15025.9321 | 2 | 7512.96603 | 0.48205245 | 0.62081147 |
| Error | 670170.938 | 43 | 15585.3706 |  |  |
| Total | 685196.87 | 45 |  |  |  |

Reward Rate
| Source | SS | df | MS | F | Prob>F |
| --- | --- | --- | --- | --- | --- |
| Power | 5.14E-05 | 2 | 2.57E-05 | 0.00906747 | 0.99097541 |
| Error | 0.12192428 | 43 | 0.00283545 |  |  |
| Total | 0.1219757 | 45 |  |  |  |

**Supplementary Table 6-3.** Statistical results for the effect of region and stimulation on probability to switch, related to Fig. 6I.

**Dbh-Cre;Ai32**
| Source | Sum Sq. | df | Singular? | Mean Sq. | F | Prob>F |
| --- | --- | --- | --- | --- | --- | --- |
| Power | 0.00287031 | 1 | 0 | 0.00287031 | 5.61667048 | 0.02270168 |
| Region | 0.00617679 | 3 | 0 | 0.00205893 | 4.02895583 | 0.01353153 |
| power*region | 0.000573 | 3 | 0 | 0.000191 | 0.3737537 | 0.77237318 |
| Error | 0.02044133 | 40 | 0 | 0.00051103 |  |  |
| Total | 0.03006143 | 47 | 0 |  |  |  |

Post-Hoc for region
| Group A | Group B | Lower Limit | A-B | Upper Limit | P-value |
| --- | --- | --- | --- | --- | --- |
| region=M2-L | region=M2-R | -0.0405306 | -0.0157933 | 0.00894397 | 0.33137496 |
| region=M2-L | region=V1-L | -0.0490326 | -0.0242953 | 0.00044197 | 0.05584813 |
| region=M2-L | region=V1-R | -0.0548628 | -0.0301256 | -0.0053883 | 0.01160543 |
| region=M2-R | region=V1-L | -0.0332393 | -0.008502 | 0.01623526 | 0.79367161 |
| region=M2-R | region=V1-R | -0.0390695 | -0.0143323 | 0.01040498 | 0.41647849 |
| region=V1-L | region=V1-R | -0.0305675 | -0.0058303 | 0.01890699 | 0.92119217 |

Post-Hoc for power
| Group A | Group B | Lower Limit | A-B | Upper Limit | P-value |
| --- | --- | --- | --- | --- | --- |
| Power=0mW | Power=3mW | -0.0304981 | -0.0168194 | -0.0031406 | 0.01723266 |

**ChAt-Cre;Ai32**
| Source | Sum Sq. | df | Singular? | Mean Sq. | F | Prob>F |
| --- | --- | --- | --- | --- | --- | --- |
| stim | 4.60E-05 | 1 | 0 | 4.60E-05 | 0.06706715 | 0.79676327 |
| region | 0.00204866 | 3 | 0 | 0.00068289 | 0.99466637 | 0.40338684 |
| stim*region | 0.00115308 | 3 | 0 | 0.00038436 | 0.55984416 | 0.64411262 |
| Error | 0.03295426 | 48 | 0 | 0.00068655 |  |  |
| Total | 0.03620204 | 55 | 0 |  |  |  |

**Supplementary Table 6-4.** Statistical results for the effect of region on overall session, related to Fig. 6J.

| Source | SS | df | MS | F | Prob>F |
| --- | --- | --- | --- | --- | --- |
| Region | 0.18972055 | 3 | 0.06324018 | 2.48832741 | 0.06710368 |
| Error | 1.85527574 | 73 | 0.02541474 |  |  |
| Total | 2.0449963 | 76 |  |  |  |

Number of Trials
| Source | SS | df | MS | F | Prob>F |
| --- | --- | --- | --- | --- | --- |
| Region | 29323.8377 | 3 | 9774.61255 | 0.67280948 | 0.57131416 |
| Error | 1118660.16 | 77 | 14528.0541 |  |  |
| Total | 1147984 | 80 |  |  |  |

Reward Rate
| Source | SS | df | MS | F | Prob>F |
| --- | --- | --- | --- | --- | --- |
| Region | 0.04258502 | 3 | 0.01419501 | 2.64214867 | 0.05522146 |
| Error | 0.41368431 | 77 | 0.00537252 |  |  |
| Total | 0.45626932 | 80 |  |  |  |

**ChAt-Cre;Ai32** Entropy
| Source | SS | df | MS | F | Prob>F |
| --- | --- | --- | --- | --- | --- |
| Region | 0.01488469 | 3 | 0.00496156 | 0.32650571 | 0.80619028 |
| Error | 1.77792625 | 117 | 0.01519595 |  |  |
| Total | 1.79281094 | 120 |  |  |  |

Number of Trials
| Source | SS | df | MS | F | Prob>F |
| --- | --- | --- | --- | --- | --- |
| Region | 945.888715 | 3 | 315.296238 | 0.02759786 | 0.99378347 |
| Error | 1336685.63 | 117 | 11424.6635 |  |  |
| Total | 1337631.52 | 120 |  |  |  |

Reward Rate
| Source | SS | df | MS | F | Prob>F |
| --- | --- | --- | --- | --- | --- |
| Region | 0.00780693 | 3 | 0.00260231 | 1.23293022 | 0.30096393 |
| Error | 0.24694836 | 117 | 0.00211067 |  |  |
| Total | 0.25475529 | 120 |  |  |  |

Post-Hoc for Reward Rate
| Group A | Group B | Lower Limit | A-B | Upper Limit | P-value |
| --- | --- | --- | --- | --- | --- |
| M2-L | M2-R | -0.057629 | -0.0180601 | 0.0215088 | 0.64422361 |
| M2-L | V1-L | -0.0588981 | -0.0193292 | 0.02023972 | 0.59174233 |
| M2-L | V1-R | -0.030134 | 0.0079956 | 0.04612516 | 0.94957223 |
| M2-R | V1-L | -0.040838 | -0.0012691 | 0.03829984 | 0.99979959 |
| M2-R | V1-R | -0.0120738 | 0.02605572 | 0.06418528 | 0.29505019 |
| V1-L | V1-R | -0.0108048 | 0.0273248 | 0.06545436 | 0.25405443 |

## Reference

Ali F, Kwan AC (2020) Interpreting in vivo calcium signals from neuronal cell bodies, axons, and dendrites: a review. Neurophotonics 7:011402.

Amaral DG, Sinnamon HM (1977) The locus coeruleus: neurobiology of a central noradrenergic nucleus. Prog Neurobiol 9:147–196.

Arnsten A, Goldman-Rakic PS (1985) Alpha 2-adrenergic mechanisms in prefrontal cortex associated with cognitive decline in aged nonhuman primates. Science (New York, NY) 230:1273–1276.

Arnsten AF, Wang MJ, Paspalas CD (2012) Neuromodulation of thought: flexibilities and vulnerabilities in prefrontal cortical network synapses. Neuron 76:223–239.

Aston-Jones G, Cohen JD (2005) An integrative theory of locus coeruleus-norepinephrine function: adaptive gain and optimal performance. Annu Rev Neurosci 28:403–450.

Aston-Jones G, Rajkowski J, Cohen J (1999) Role of locus coeruleus in attention and behavioral flexibility. Biological psychiatry 46:1309–1320.

Atilgan H, Murphy CE, Wang H, Ortega HK, Pinto L, Kwan AC (2022) Change point estimation by the mouse medial frontal cortex during probabilistic reward learning. bioRxiv.

Azcorra M, Gaertner Z, Davidson C, He Q, Kim H, Nagappan S, Hayes CK, Ramakrishnan C, Fenno L, Kim YS, Deisseroth K, Longnecker R, Awatramani R, Dombeck DA (2023) Unique functional responses differentially map onto genetic subtypes of dopamine neurons. Nat Neurosci 26:1762–1774.

Barraclough DJ, Conroy ML, Lee D (2004) Prefrontal cortex and decision making in a mixed-strategy game. Nat Neurosci 7:404–410.

Barthas F, Kwan AC (2017) Secondary Motor Cortex: Where ’Sensory’ Meets ’Motor’ in the Rodent Frontal Cortex. Trends Neurosci 40:181–193.

Behrens TE, Woolrich MW, Walton ME, Rushworth MF (2007) Learning the value of information in an uncertain world. Nat Neurosci 10:1214–1221.

Berridge CW, Waterhouse BD (2003) The locus coeruleus–noradrenergic system: modulation of behavioral state and state-dependent cognitive processes. Brain Research Reviews 42:33–84.

Bouret S, Sara SJ (2005) Network reset: a simplified overarching theory of locus coeruleus noradrenaline function. Trends Neurosci 28:574–582.

Bouret S, Richmond BJ (2015) Sensitivity of locus ceruleus neurons to reward value for goal-directed actions. J Neurosci 35:4005–4014.

Buzsaki G, Bickford RG, Ponomareff G, Thal LJ, Mandel R, Gage FH (1988) Nucleus basalis and thalamic control of neocortical activity in the freely moving rat. J Neurosci 8:4007–4026.

Chudasama Y, Dalley JW, Nathwani F, Bouger P, Robbins TW (2004) Cholinergic modulation of visual attention and working memory: dissociable effects of basal forebrain 192-IgG-saporin lesions and intraprefrontal infusions of scopolamine. Learn Mem 11:78–86.

Clayton EC, Rajkowski J, Cohen JD, Aston-Jones G (2004) Phasic activation of monkey locus ceruleus neurons by simple decisions in a forced-choice task. J Neurosci 24:9914–9920.

Croxson PL, Kyriazis DA, Baxter MG (2011) Cholinergic modulation of a specific memory function of prefrontal cortex. Nat Neurosci 14:1510–1512.

Dalley JW, McGaughy J, O’Connell MT, Cardinal RN, Levita L, Robbins TW (2001) Distinct changes in cortical acetylcholine and noradrenaline efflux during contingent and noncontingent performance of a visual attentional task. J Neurosci 21:4908–4914.

Dayan P, Yu AJ (2006) Phasic norepinephrine: a neural interrupt signal for unexpected events. Network 17:335–350.

de Gee JW, Knapen T, Donner TH (2014) Decision-related pupil dilation reflects upcoming choice and individual bias. Proc Natl Acad Sci U S A 111:E618–625.

Descarries L, Lapierre Y (1973) Noradrenergic axon terminals in the cerebral cortex of rat. I. Radioautographic visualization after topical application of DL-[3H]Norepinephrine. Brain Res 51:141–160.

Doya K (2002) Metalearning and neuromodulation. Neural Netw 15:495–506.

Einhauser W, Koch C, Carter OL (2010) Pupil dilation betrays the timing of decisions. Front Hum Neurosci 4:18.

Eldar E, Cohen JD, Niv Y (2013) The effects of neural gain on attention and learning. Nat Neurosci 16:1146–1153.

Everitt BJ, Robbins TW (1997) Central cholinergic systems and cognition. Annu Rev Psychol 48:649–684.

Feng J, Dong H, Lischinsky J, Zhou J, Deng F, Zhuang C, Miao X, Wang H, Xie H, Cui G, Lin D, Li Y (2023) Monitoring norepinephrine release in vivo using next-generation GRABNE sensors. bioRxiv.

Feng J, Zhang C, Lischinsky JE, Jing M, Zhou J, Wang H, Zhang Y, Dong A, Wu Z, Wu H, Chen W, Zhang P, Zou J, Hires SA, Zhu JJ, Cui G, Lin D, Du J, Li Y (2019) A Genetically Encoded Fluorescent Sensor for Rapid and Specific In Vivo Detection of Norepinephrine. Neuron 102:745–761 e748.

Froemke RC, Merzenich MM, Schreiner CE (2007) A synaptic memory trace for cortical receptive field plasticity. Nature 450:425–429.

Goard M, Dan Y (2009) Basal forebrain activation enhances cortical coding of natural scenes. Nat Neurosci 12:1444–1449.

Gritton HJ, Howe WM, Mallory CS, Hetrick VL, Berke JD, Sarter M (2016) Cortical cholinergic signaling controls the detection of cues. Proc Natl Acad Sci U S A 113:E1089–1097.

Hangya B, Ranade SP, Lorenc M, Kepecs A (2015) Central Cholinergic Neurons Are Rapidly Recruited by Reinforcement Feedback. Cell 162:1155–1168.

Hasselmo ME, Bower JM (1993) Acetylcholine and memory. Trends Neurosci 16:218–222.

Henny P, Jones BE (2008) Projections from basal forebrain to prefrontal cortex comprise cholinergic, GABAergic and glutamatergic inputs to pyramidal cells or interneurons. Eur J Neurosci 27:654–670.

Herrero JL, Roberts MJ, Delicato LS, Gieselmann MA, Dayan P, Thiele A (2008) Acetylcholine contributes through muscarinic receptors to attentional modulation in V1. Nature 454:1110–1114.

Howe MW, Dombeck DA (2016) Rapid signalling in distinct dopaminergic axons during locomotion and reward. Nature 535:505–510.

Jing M et al. (2020) An optimized acetylcholine sensor for monitoring in vivo cholinergic activity. Nat Methods 17:1139–1146.

Jouvet M (1969) Biogenic amines and the states of sleep. Science (New York, NY) 163:32–41.

Kagiampaki Z et al. (2023) Sensitive multicolor indicators for monitoring norepinephrine in vivo. Nat Methods 20:1426–1436.

Kilgard MP (1998) Cortical Map Reorganization Enabled by Nucleus Basalis Activity. Science 279:1714–1718.

Kim JH, Jung AH, Jeong D, Choi I, Kim K, Shin S, Kim SJ, Lee SH (2016) Selectivity of Neuromodulatory Projections from the Basal Forebrain and Locus Ceruleus to Primary Sensory Cortices. J Neurosci 36:5314–5327.

Laszlovszky T, Schlingloff D, Hegedus P, Freund TF, Gulyas A, Kepecs A, Hangya B (2020) Distinct synchronization, cortical coupling and behavioral function of two basal forebrain cholinergic neuron types. Nat Neurosci 23:992–1003.

Lee D (2008) Game theory and neural basis of social decision making. Nat Neurosci 11:404–409.

Lee D, Conroy ML, McGreevy BP, Barraclough DJ (2004) Reinforcement learning and decision making in monkeys during a competitive game. Brain Res Cogn Brain Res 22:45–58.

Lin SC, Nicolelis MA (2008) Neuronal ensemble bursting in the basal forebrain encodes salience irrespective of valence. Neuron 59:138–149.

Liu CH, Coleman JE, Davoudi H, Zhang K, Hussain Shuler MG (2015) Selective activation of a putative reinforcement signal conditions cued interval timing in primary visual cortex. Curr Biol 25:1551–1561.

Lohani S, Moberly AH, Benisty H, Landa B, Jing M, Li Y, Higley MJ, Cardin JA (2022) Spatiotemporally heterogeneous coordination of cholinergic and neocortical activity. Nat Neurosci 25:1706–1713.

Loughlin SE, Foote SL, Fallon JH (1982) Locus coeruleus projections to cortex: topography, morphology and collateralization. Brain Res Bull 9:287–294.

Madisen L et al. (2012) A toolbox of Cre-dependent optogenetic transgenic mice for light-induced activation and silencing. Nat Neurosci 15:793–802.

Maho C, Hars B, Edeline J-M, Hennevin E (1995) Conditioned changes in the basal forebrain: Relations with learning-induced cortical plasticity. Psychobiology 23:10–25.

Martins AR, Froemke RC (2015) Coordinated forms of noradrenergic plasticity in the locus coeruleus and primary auditory cortex. Nat Neurosci 18:1483–1492.

Mathis A, Mamidanna P, Cury KM, Abe T, Murthy VN, Mathis MW, Bethge M (2018) DeepLabCut: markerless pose estimation of user-defined body parts with deep learning. Nat Neurosci 21:1281–1289.

McCormick DA, Pape HC, Williamson A (1991) Actions of norepinephrine in the cerebral cortex and thalamus: implications for function of the central noradrenergic system. Progress in brain research 88:293–305.

McGaughy J, Dalley JW, Morrison CH, Everitt BJ, Robbins TW (2002) Selective behavioral and neurochemical effects of cholinergic lesions produced by intrabasalis infusions of 192 IgG-saporin on attentional performance in a five-choice serial reaction time task. J Neurosci 22:1905–1913.

Moore R, Bloom F (1979) Central catecholamine neuron systems: anatomy and physiology of the norepinephrine and epinephrine systems. Annual review of neuroscience 2:113–168.

Nath T, Mathis A, Chen AC, Patel A, Bethge M, Mathis MW (2019) Using DeepLabCut for 3D markerless pose estimation across species and behaviors. Nat Protoc 14:2152–2176.

Parikh V, Kozak R, Martinez V, Sarter M (2007) Prefrontal acetylcholine release controls cue detection on multiple timescales. Neuron 56:141–154.

Patriarchi T, Mohebi A, Sun J, Marley A, Liang R, Dong C, Puhger K, Mizuno GO, Davis CM, Wiltgen B, von Zastrow M, Berke JD, Tian L (2020) An expanded palette of dopamine sensors for multiplex imaging in vivo. Nat Methods 17:1147–1155.

Picciotto MR, Higley MJ, Mineur YS (2012) Acetylcholine as a neuromodulator: cholinergic signaling shapes nervous system function and behavior. Neuron 76:116–129.

Pinto L, Rajan K, DePasquale B, Thiberge SY, Tank DW, Brody CD (2019) Task-Dependent Changes in the Large-Scale Dynamics and Necessity of Cortical Regions. Neuron 104:810–824 e819.

Pinto L, Goard MJ, Estandian D, Xu M, Kwan AC, Lee SH, Harrison TC, Feng G, Dan Y (2013) Fast modulation of visual perception by basal forebrain cholinergic neurons. Nat Neurosci 16:1857–1863.

Pnevmatikakis EA, Giovannucci A (2017) NoRMCorre: An online algorithm for piecewise rigid motion correction of calcium imaging data. Journal of neuroscience methods 291:83–94.

Reimer J, McGinley MJ, Liu Y, Rodenkirch C, Wang Q, McCormick DA, Tolias AS (2016) Pupil fluctuations track rapid changes in adrenergic and cholinergic activity in cortex. Nat Commun 7:13289.

Rossi J, Balthasar N, Olson D, Scott M, Berglund E, Lee CE, Choi MJ, Lauzon D, Lowell BB, Elmquist JK (2011) Melanocortin-4 receptors expressed by cholinergic neurons regulate energy balance and glucose homeostasis. Cell Metab 13:195–204.

Saper CB (1984) Organization of cerebral cortical afferent systems in the rat. II. Magnocellular basal nucleus. J Comp Neurol 222:313–342.

Sara SJ (2009) The locus coeruleus and noradrenergic modulation of cognition. Nat Rev Neurosci 10:211–223.

Schwarz LA, Miyamichi K, Gao XJ, Beier KT, Weissbourd B, DeLoach KE, Ren J, Ibanes S, Malenka RC, Kremer EJ, Luo L (2015) Viral-genetic tracing of the input-output organization of a central noradrenaline circuit. Nature 524:88–92.

Servan-Schreiber D, Printz H, Cohen JD (1990) A network model of catecholamine effects: gain, signal-to-noise ratio, and behavior. Science (New York, NY) 249:892–895.

Siniscalchi MJ, Wang H, Kwan AC (2019) Enhanced Population Coding for Rewarded Choices in the Medial Frontal Cortex of the Mouse. Cereb Cortex.

Siniscalchi MJ, Phoumthipphavong V, Ali F, Lozano M, Kwan AC (2016) Fast and slow transitions in frontal ensemble activity during flexible sensorimotor behavior. Nat Neurosci 19:1234–1242.

Starkweather CK, Gershman SJ, Uchida N (2018) The Medial Prefrontal Cortex Shapes Dopamine Reward Prediction Errors under State Uncertainty. Neuron 98:616–629 e616.

Sturgill J, Hegedus P, Li S, Chevy Q, Siebels A, Jing M, Li Y, Hangya B, Kepecs A (2020) Basal forebrain-derived acetylcholine encodes valence-free reinforcement prediction error. bioRxiv.

Su Z, Cohen JY (2022) Two types of locus coeruleus norepinephrine neurons drive reinforcement learning. biorxiv.

Swanson L, Hartman B (1975) The central adrenergic system. An immunofluorescence study of the location of cell bodies and their efferent connections in the rat utilizing dopamine-B-hydroxylase as a marker. Journal of Comparative Neurology 163:467–505.

Teles-Grilo Ruivo LM, Baker KL, Conway MW, Kinsley PJ, Gilmour G, Phillips KG, Isaac JTR, Lowry JP, Mellor JR (2017) Coordinated Acetylcholine Release in Prefrontal Cortex and Hippocampus Is Associated with Arousal and Reward on Distinct Timescales. Cell Rep 18:905–917.

Tervo DG, Proskurin M, Manakov M, Kabra M, Vollmer A, Branson K, Karpova AY (2014) Behavioral variability through stochastic choice and its gating by anterior cingulate cortex. Cell 159:21–32.

Tervo DGR, Kuleshova E, Manakov M, Proskurin M, Karlsson M, Lustig A, Behnam R, Karpova AY (2021) The anterior cingulate cortex directs exploration of alternative strategies. Neuron 109:1876–1887 e1876.

Tillage RP, Sciolino NR, Plummer NW, Lustberg D, Liles LC, Hsiang M, Powell JM, Smith KG, Jensen P, Weinshenker D (2020) Elimination of galanin synthesis in noradrenergic neurons reduces galanin in select brain areas and promotes active coping behaviors. Brain Struct Funct 225:785–803.

Totah NK, Neves RM, Panzeri S, Logothetis NK, Eschenko O (2018) The Locus Coeruleus Is a Complex and Differentiated Neuromodulatory System. Neuron 99:1055–1068 e1056.

Uematsu A, Tan BZ, Ycu EA, Cuevas JS, Koivumaa J, Junyent F, Kremer EJ, Witten IB, Deisseroth K, Johansen JP (2017) Modular organization of the brainstem noradrenaline system coordinates opposing learning states. Nat Neurosci 20:1602–1611.

van Holstein M, Floresco SB (2020) Dissociable roles for the ventral and dorsal medial prefrontal cortex in cue-guided risk/reward decision making. Neuropsychopharmacology 45:683–693.

Van Slooten JC, Jahfari S, Knapen T, Theeuwes J (2018) How pupil responses track value-based decision-making during and after reinforcement learning. PLoS Comput Biol 14:e1006632.

Vertechi P, Lottem E, Sarra D, Godinho B, Treves I, Quendera T, Oude Lohuis MN, Mainen ZF (2020) Inference-Based Decisions in a Hidden State Foraging Task: Differential Contributions of Prefrontal Cortical Areas. Neuron 106:166–176.e166.

Wang H, Kwan AC (2023) Competitive and cooperative games for probing the neural basis of social decision-making in animals. Neurosci Biobehav Rev 149:105158.

Wang H, Ortega HK, Atilgan H, Murphy CE, Kwan AC (2022) Pupil correlates of decision variables in mice playing a competitive mixed-strategy game. Eneuro 9.

Yang JH, Kwan AC (2021) Secondary motor cortex: Broadcasting and biasing animal’s decisions through long-range circuits. Int Rev Neurobiol 158:443–470.

Yu AJ, Dayan P (2002) Acetylcholine in cortical inference. Neural Netw 15:719–730.

Zaborszky L, Csordas A, Mosca K, Kim J, Gielow MR, Vadasz C, Nadasdy Z (2015) Neurons in the basal forebrain project to the cortex in a complex topographic organization that reflects corticocortical connectivity patterns: an experimental study based on retrograde tracing and 3D reconstruction. Cereb Cortex 25:118–137.

Zhu PK, Zheng WS, Zhang P, Jing M, Borden PM, Ali F, Guo K, Feng J, Marvin JS, Wang Y, Wan J, Gan L, Kwan AC, Lin L, Looger LL, Li Y, Zhang Y (2020) Nanoscopic Visualization of Restricted Nonvolume Cholinergic and Monoaminergic Transmission with Genetically Encoded Sensors. Nano Lett 20:4073–4083.

Zhuo Y et al. (2023) Improved green and red GRAB sensors for monitoring dopaminergic activity in vivo. Nat Methods.

